# The Effects of Bipolar Disorder Granule Cell Hyperexcitability and Lithium Therapy on Pattern Separation in a Computational Model of the Dentate Gyrus

**DOI:** 10.1101/2024.04.09.588764

**Authors:** Selena Singh, Anouar Khayachi, Shani Stern, Thomas Trappenberg, Martin Alda, Abraham Nunes

## Abstract

Induced pluripotent stem cell (iPSC) derived hippocampal dentate granule cell-like neurons from individuals with bipolar disorder (BD) are hyperexcitable and more spontaneously active relative to healthy control (HC) neurons. These abnormalities are normalised after the application of lithium in neurons derived from lithium responders (LR) only. How these abnormalities impact hippocampal microcircuit computation is not understood. We aimed to investigate the impacts of BD-associated abnormal granule cell (GC) activity on pattern separation (PS) using a computational model of the dentate gyrus (DG). We used parameter optimization to fit the parameters of biophysically realistic granule cell (GC) models to electrophysiological data from iPSC GCs from patients with BD. These cellular models were incorporated into DG networks to assess impacts on PS using an adapted spatiotemporal task. Relationships between BD, lithium and spontaneous activity were analysed using linear mixed effects modelling. Lithium and BD negatively impacted PS, consistent with clinical reports of cognitive slowing and memory impairment during lithium therapy. By normalising spontaneous activity levels, lithium improved PS performance in LRs only. Improvements in PS after lithium therapy in LRs may therefore be attributable to the normalisation of spontaneous activity levels, rather than reductions in GC intrinsic excitability as we hypothesised. Our results agree with a hypothesised relationship between behavioural mnemonic discrimination and DG PS, as previous research has suggested that mnemonic discrimination improves after lithium therapy in lithium responders only. Our work can be expanded on in the future by simulating the effects of lithium-induced neurogenesis on PS.

## 1 INTRODUCTION

Bipolar disorder (BD) is a mood disorder with unknown aetiology characterised by recurrent episodes of mania and depression [1], as well as cognitive impairments that are functionally impactful [2, 3] and persist during euthymia [4]. Both encoding and retrieval processes for verbal material are impacted [5–7], and additional studies link BD with poor autobiographical memory specificity [8–10] and recognition memory deficits [11, 12]. Identifying the neural mechanisms of neurocognitive impairment may inform treatment development and reduce disease burden.

The hippocampus is important for both memory and emotion, and may be involved in the pathogenesis of BD. The hippocampus is critical for encoding complex associative and autobiographical memories [13–15], and is ideally suited to promote contextually appropriate responses because of its connectivity with brain systems involved in executive functioning, motivation, stress response, and emotion [16, 17]. Recent theories have proposed that temporal context-dependent representations created by the integration of amygdalar and prefrontal inputs by the hippocampus may constrain emotional responses to their appropriate contexts, which may protect against psychopathology [17]. Disruptions in hippocampal function will therefore not only impact memory processes, but also have downstream impacts on emotion and cognition by influencing dynamics within cortico-limbic-subcortical circuits, and this dynamic role of the hippocampus has been proposed to play a large role in BD pathogenesis [18]. Indeed, anatomical, functional imaging, and physiological hippocampal abnormalities have been reported in BD, and are reviewed briefly below.

Hippocampal abnormalities in BD include reduced hippocampal volume, reduced inhibitory interneuron expression, and an increase in recurrent excitatory projections between dentate granule cells as reported in post-mortem studies [19– 21] (for review, see [18]). A meta-analysis of functional magnetic resonance imaging studies has reported hyperactivity in limbic (i.e., parahippocampal, hippocampal and amygdalar) areas in BD relative to healthy individuals [22]. Lithium, the gold-standard prophylactic for BD, may protect against BD-associated hippocampal volume loss [23, 24]. In summary, these studies demonstrate that the hippocampus is a brain area of interest in BD, but do not describe the physiological abnormalities that would impact neural computation, leading to the cognitive deficits described earlier.

Induced pluripotent stem cell (iPSC) technology has been recently used to create hippocampal cell models *in vitro* from stem cells derived from individuals with BD, to facilitate study of the cellular physiological abnormalities of BD [25, 26]. Lithium responsive and non-responsive BD iPSC models of the pyramidal CA3 neuron and dentate granule cell (GC) have been created to date, and these neurons indeed have abnormal physiological properties that differ between models derived from lithium responders (LR) and lithium non-responders (NR)[25–28]. Both LR and NR iPSC neurons are hyperexcitable relative to healthy controls, and this hyperexcitability is normalised after application of lithium only for neurons derived from LRs [26]. LR-BD cell models also demonstrate elevated spontaneous activity levels relative to NR-BD and healthy control models; lithium normalises spontaneous activity levels in LR neurons as well as hyperexcitability [25]. In other words, response to lithium at the cellular level corresponds to the patient’s clinical response to lithium, suggesting that this cellular phenomenon may be a useful biomarker of treatment response in BD. These neurons may also play a core role in BD’s pathophysiology, and explain lithium’s mechanism of action. How these abnormalities impact hippocampal microcircuit neural computation, contributing to BD-associated cognitive and memory impairments, is not yet understood.

The present study aims to investigate the impacts of GC hyperexcitability in lithium responsive and nonresponsive BD on the neural computation called pattern separation (PS) widely attributed to the hippocampal dentate gyrus (DG). PS is a computation that involves mapping highly overlapping and similar inputs onto less overlapping and dissimilar outputs [29], aiding the hippocampus with encoding precise memories with minimal interference. The DG is ideally suited to perform this computation due to the sparse, competitive firing of mature GCs that are tightly controlled by powerful inhibitory interneurons [30–32]. Evidence from rodent electrophysiology supports the idea of the DG performing PS by demonstrating that there is less correlated activity in the DG than in the entorhinal cortex and CA3 in response to slight changes in environmental stimuli [33, 34]. GCs themselves also have been shown to produce separated representations of inputs by shifting output spike times [35, 36], highlighting the importance of GC physiology for PS. PS within the DG may also be behaviourally relevant, as it has been hypothesised to underlie performance on high-interference memory tasks, such as mnemonic discrimination (MD) [37], which is a phenomenon that involves discerning between stimuli with highly overlapping qualitative properties [38]. Following this hypothesis, DG PS deficits may therefore serve as predictors for deficits in performance on high-interference memory tasks, such as MD, in BD. Interestingly, results from a pilot study of MD performance in BD have suggested that lithium may improve MD in LRs only [39].

This work aims to predict what the consequences of GC hyperexcitability on DG PS are, building a bridge between *in vitro* iPSC [26, 27, 40] and *in vivo* behavioural work [39]. We hypothesise that BD-specific GC hyperexcitability will lead to PS impairments, which will resolve after the normalisation of hyperexcitability via application of lithium in LRs. We test this hypothesis by integrating detailed biophysical models of these abnormal BD GCs into a larger DG network model, and evaluate the network’s PS abilities. Our study will help elucidate neural computations underlying some of the cognitive and memory-related impairments in BD.

## 2 METHODS

To study the effects of LR and NR GC hyperexcitability, spontaneous activity, and effects of lithium on PS, we developed biophysically realistic computational models based on empirical data from patient-derived iPSC neurons. We outline details of our approach in the Supplementary Materials, and present an intuitive description here. A schematic walk-through of our methods is shown in Figure 1.

**Figure 1:**
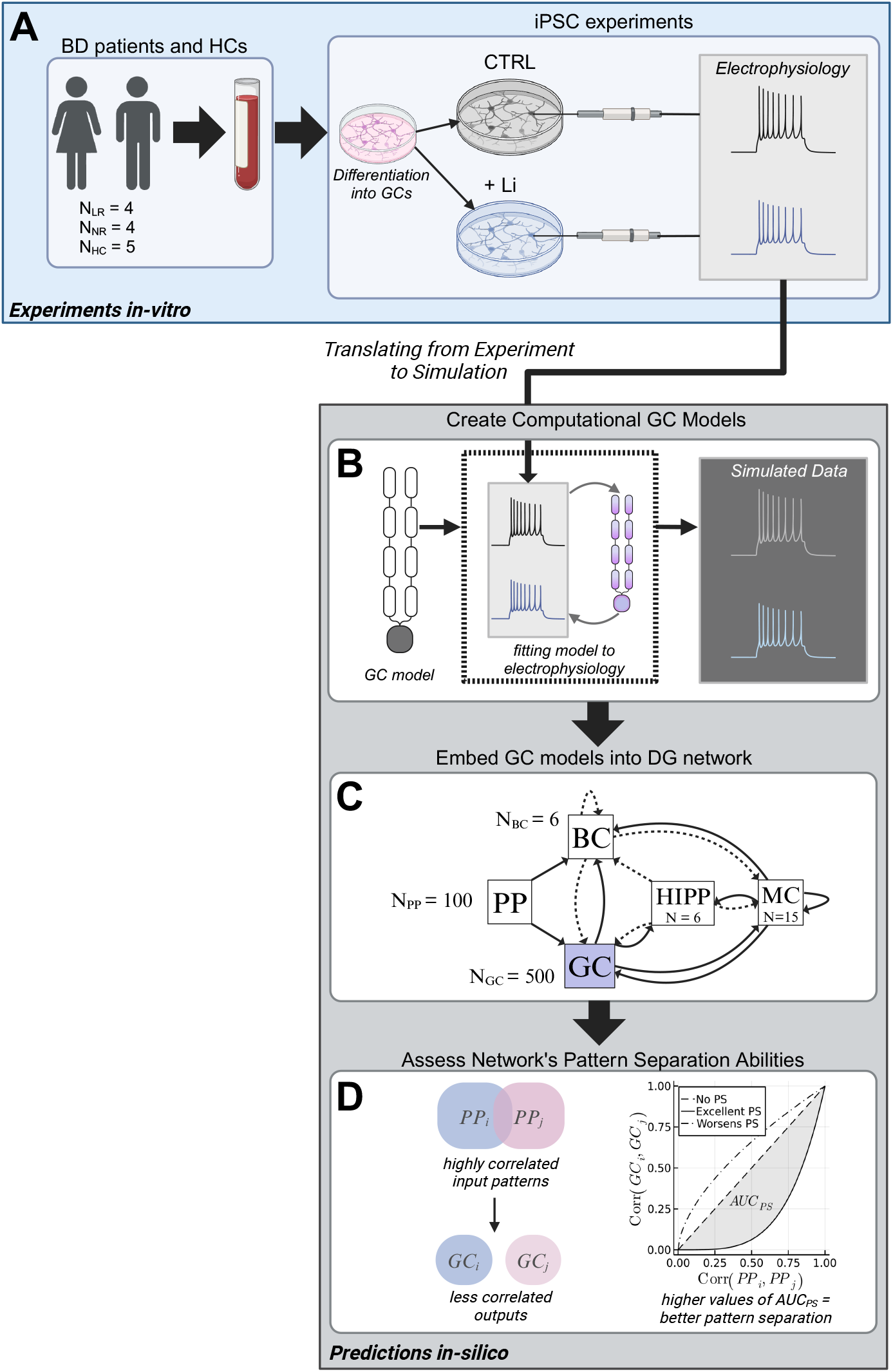
Schematic of study methodology: from human participants to computational modelling. A) Blood samples were first collected from individuals with bipolar disorder (BD) (both lithium responders, LRs, and non-responders, NRs) and healthy controls (HCs), and cells were reprogrammed into granule cell (GC)-like neurons. Half of the GCs were exposed to lithium, and the electrophysiological properties of these neurons were studied. These results have been previously reported by Khayachi et al., 2023 [40]. B) We used these electrophysiological data (frequency-current and current-voltage curves specifically) to fit the parameters of a model GC such that the model generated the same electrophysiological behaviour as the *in vitro* GCs. Note: spike trains shown here are for illustrative purposes only, and are not real GC spike trains. C) These model GCs were then incorporated into a biophysical dentate gyrus (DG) network, to form model DGs for LRs, NRs and HCs. PP: perforant path, BC: basket cell, HIPP: hilar perforant path cells, MC: mossy cell. Solid lines indicate excitatory connections, and dashed lines indicate inhibitory connections. N indicates the number of cells per population. This circuit diagram was adapted from our previous paper [44]. D) The pattern separation (PS) performance of these networks were then assessed, by presenting the network with a series of partially overlapping PP input patterns, and assessing whether the resulting output patterns were less correlated. Plotting the correlation between pairs of input patterns and resulting output patterns against each other generated a PS curve. The area between the diagonal and this pattern separation curve (*AUC*_*P S*_) summarised the network’s PS abilities, with larger *AUC*_*P S*_ values representing better PS.

### 2.1 Developing GC Models

#### 2.1.1 Description of iPSC-derived Dentate Gyrus Granule Cell-Like Neurons

First, iPSC neurons were reprogrammed from lymphocytes and peripheral blood mononuclear cells (PBMC) taken from blood samples from consenting participants. Detailed methodology describing iPSC differentiation and cell culture protocols for the iPSC neurons used to inform our modelling have been previously described [40]. Briefly, blood samples were collected from 8 BD patients (4 LRs, and 4 NRs), and 5 healthy control (HC) participants (Figure 1A). After lymphocyte and PBMC isolation, followed by iPSC differentiation as described previously [26, 40], about half of the neurons per group were exposed to therapeutic levels of lithium (1.5 mM), for 7 days. Therefore, there were the following number of iPSC GC-like neurons per group used for whole-cell patch-clamp recordings: LR: (*n*_*Li*_= 49, *n*_*CT RL*_= 55); NR: (*n*_*Li*_ = 41, *n*_*CT RL*_= 45); HC: (*n*_*Li*_ = 42, *n*_*CT RL*_=40). Sodium and potassium current-voltage (IV), and frequency-current (FI) relationships were acquired in voltage-clamp and current-clamp modes respectively.

#### 2.1.2 Computational Model of a Dentate GC

We then adapted an established computational model of the hippocampal dentate GC [41–44] implemented in the NEURON simulation environment (v. 8.0) [45]. This model has two identical dendrites with four compartments each, and a single compartment for the soma (Figure 1B). Distributed along the somatodendritic tree are 11 different ion-channels: fast sodium (Na), fast and slow delayed rectifier potassium, A-type potassium, large conductance calcium, voltage-dependent potassium, small conductance calcium-dependent potassium, T-type, N-type, and L-type voltage-gated calcium, inward-rectifier potassium and the tonic GABAA chloride channel. The dynamics of each of these channels are described by sets of differential equations that are parameterized to produce behaviour consistent with what is observed in real-world GCs. Together, these parameters govern the intrinsic excitability and behaviour of these model neurons. To ensure these models captured the behaviour of real-world iPSC-derived neurons from BD and HC participants, we fit these parameters using numerical optimization to electrophysiological data described in Section 2.1.1 (Figure 1B).

#### 2.1.3 Numerical Optimization-Based Fitting of Computational Models to Cellular Data

Parameter optimization was done using evolutionary algorithms in the *inspyred* (v. 1.0) and *NetPyNe* (v. 1.0.0.2) Python packages. The objective function minimized was the averaged mean squared error between model and iPSC-neuron FI and IV curves, for iPSC neurons with and without exposure to lithium (“LITM” and “CTRL”, respectively). This procedure therefore generated six models: HC-CTRL, HC-LITM, LR-CTRL, LR-LITM, NR-CTRL and NR-LITM. Evolutionary algorithms perform parameter optimization by iteratively mutating, then evaluating the “fitness” of a parameter set [46]. We deemed the model fits to be satisfactory if each simulated data point fell within the empirical standard error of the mean.

#### 2.1.4 Granule Cell-Like Neuron Models for Lithium Nonresponders

Experimental data failed to show a statistically significant effect of lithium exposure on FI and IV relationships for the NR iPSC GCs, meaning these two curves were statistically identical. Therefore, to produce a NR-LITM model, we began with the fitted NR-CTRL model and modified the parameters by increasing or decreasing their values by a random value less than 2% of the original parameter’s value to introduce some noise. This approach yielded two models with slight differences that have comparable parameter values and biophysical behaviour, which we believe are good candidates for simulating NR-CTRL and NR-LITM conditions.

#### 2.1.5 Simulation of Spontaneous Activity

Randomly selected GCs within the network were equipped with Poisson spike generators, synapsed onto GC somata, that randomly produced spikes at the following rates during the simulation: HC and NRs = 0.25 Hz; LRs = 1Hz, following previous experimental reports of spontaneous activity levels [25]. The effect of lithium on spontaneous activity was captured by setting the LR spontaneous activity level back to HC levels of 0.25 Hz [25].

### 2.2 Biophysical Network Model of the Dentate Gyrus

We then incorporated the model GCs into a DG network, to assess how BD-associated GC electrophysiological abnormalities may interfere with PS functioning. We employed a previously established conductance-based biophysical model of the DG [42–44], also implemented in the NEURON simulation environment [45], and used the original model’s geometric and topological features. Our model included 500 glutamatergic GCs (as described earlier), 6 GABAergic basket cells (BCs), 15 glutamatergic mossy cells (MCs), 6 GABAergic hilar perforant path cells (HIPP), and 100 excitatory entorhinal perforant path (PP) cells (Figure 1C). As with the GCs, the other cells in the network (BCs, MCs, HIPP) were modelled as multicompartmental Hodgkin-Huxley style neurons with a soma and varying numbers of dendrites. Details regarding these neurons can be found in our Supplementary Materials. PP cells were modelled as point processes that stimulated GCs and BCs.

All biophysical properties were kept the same as in the original model [42–44]. Parameter values for the connectivity, cellular biophysics, and synaptic double-exponential functions can also be found in our Supplementary Materials, and are also described in our previous study that employed this model [44].

#### 2.2.1 Spatiotemporal PS task

We adopted a previously established spatiotemporal PS task to assess our network’s PS functioning [42, 44, 47]. This protocol involved simulating 24 partially overlapping patterns, varying smoothly in degrees of overlap, of PP inputs over a 200 ms window. We then assessed whether pairs of the resulting output pattern representations in the GC layer of our network were less correlated than the inputs by computing Pearson correlations (Figure 1D). By using the correlation of input patterns as *x* coordinates and the correlation of output representations as *y* coordinates, plotting this relationship between inputs and outputs will produce a curve that falls below the leading diagonal if the network is performing PS (Figure 1D). In other words, highly correlated inputs should be mapped onto less correlated outputs. We computed a summary PS index defined as the area between the leading diagonal and the PS curve (*AUC*_*P S*_). Higher values of *AUC*_*P S*_ indicate stronger PS by the DG network (Figure 1D).

### 2.3 Statistical Analysis

The predicted impacts of group (HC, NR, LR), treatment (with lithium and without) and spontaneous activity (baseline vs. pathological) on standardised (i.e., z-scored) PS scores (*AUC*_*P S*_) were characterised by the following linear mixed effects model, presented here in R syntax for the *lme4* package in the R programming language [48]:

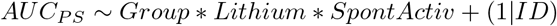

where ID refers to each simulation run, which is initialised with a different random seed to incorporate variability in network connectivity and which GCs are spontaneously active. *A priori* power calculations using the *simR* package in R showed that 14 simulation runs per experimental condition offered 80% power to detect a 5% change in *AUC*_*P S*_ at a statistical significance threshold of *α*=0.05 for the three-way interaction. Model coefficients are reported as standardised effects, in the number of standard deviations of *AUC*_*P S*_.

### 2.4 Assessing k winner-take-all dynamics

One strategy proposed to promote PS in the DG is the competitive activation of GCs facilitated by BC lateral inhibition. In our network, GCs are organised into lamellar clusters defined by BC number, with BCs projecting onto 100 GCs each. Under a winner-take-all (WTA) paradigm, the stimulation of a few GCs should lead to the inhibition of the other GCs within each lamella via the BC projections. This will promote the selection of only a few GCs to fire, or a sparse coding regime, supporting PS [32, 49–53]. To further understand the effects of BD and lithium on PS, we analysed our network’s winner-take-all dynamics as follows. We treated each spontaneous stimulation event (described in section 2.1.4) as a randomised trial, and assessed the aggregate population-level behaviour of the GCs directly stimulated (or directly activated, “AC”), and those that were not stimulated (or remaining, “RM”), within a 10 ms window post-stimulation, one lamella at a time. This population activity was averaged across 14 simulation runs. Under a WTA paradigm, we expect a peak in activity within GCs that are directly activated (these neurons are therefore the “winners”), and little to no change in behaviour within GCs that are not directly activated, suggesting tight inhibitory control, and thus a strict “selection” process, of the neurons within these lamellar microcircuits.

### 2.5 Cellular response to negative currents

Given that a WTA mechanism for PS is dependent on the inhibition of GCs, studying GC neuronal sensitivity to inhibition is essential. For this protocol, current was injected into the somatic compartment for 1s from −33pA to 0pA in 3pA steps, and the overall membrane potential was recorded, mirroring the protocol used for the iPSC GCs *in vitro*.

## 3 RESULTS

### 3.1 Fitting GC model to iPSC data

Results from our parameter fitting procedure are shown in Supplementary Figure 1, with spike trains for each model shown in Supplementary Figure 2. Lithium reduces GC hyperexcitability in LRs, but not NRs (Figure 2A). Sodium and potassium current magnitudes are also reduced after lithium administration in LRs (Figure 2B and C). Lithium also reduces excitability and sodium and potassium conductances in HCs (Figure 2). Resulting parameter values after model fitting can be found in Supplementary Materials, Table 4.

**Figure 2:**
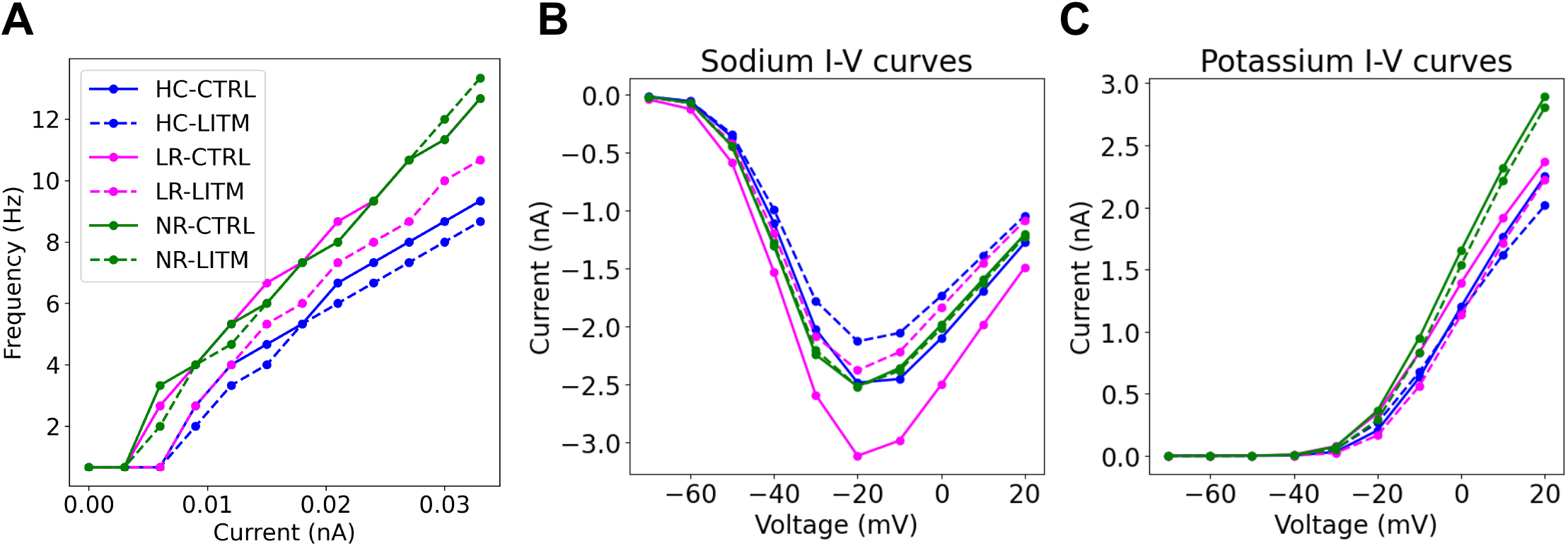
Model GC electrophysiology for HC, LR and NR models, with and without lithium. A) frequency-current relationships. BD models are more excitable than HC models, and excitability is reduced for LRs and HCs after lithium exposure. B) Sodium current-voltage relationships. Lithium reduces sodium currents for LRs and HCs. C) Potassium current-voltage relationships. NRs have greater potassium currents than LRs and HCs.

### 3.2 Effects of BD hyperexcitability and lithium on PS

The resulting PS performance for DG networks with HC and BD GC models, for baseline and pathological levels of spontaneous activity, are shown in Figure 3, and results from fitting the linear mixed effects model to the simulated data are shown in Supplementary Table 1. BD DG models performed significantly poorer PS than HC models, regardless of spontaneous activity levels and lithium treatment (LR: *β*= −2.37, CI: −2.46 – −2.29, p<0.001; NR: *β* = −0.75, CI: −0.83 – −0.66, p<0.001) (Figure 3). Lithium, independent of Group or Spontaneous Activity, significantly and negatively impacted PS in general (*β* = −1.32, CI: −1.41 – −1.24, p<0.001) (Figure 3). Although lithium reduced the excitability of HC GCs (Figure 2A), it also reduced HC PS performance (Figure 3A).

**Figure 3:**
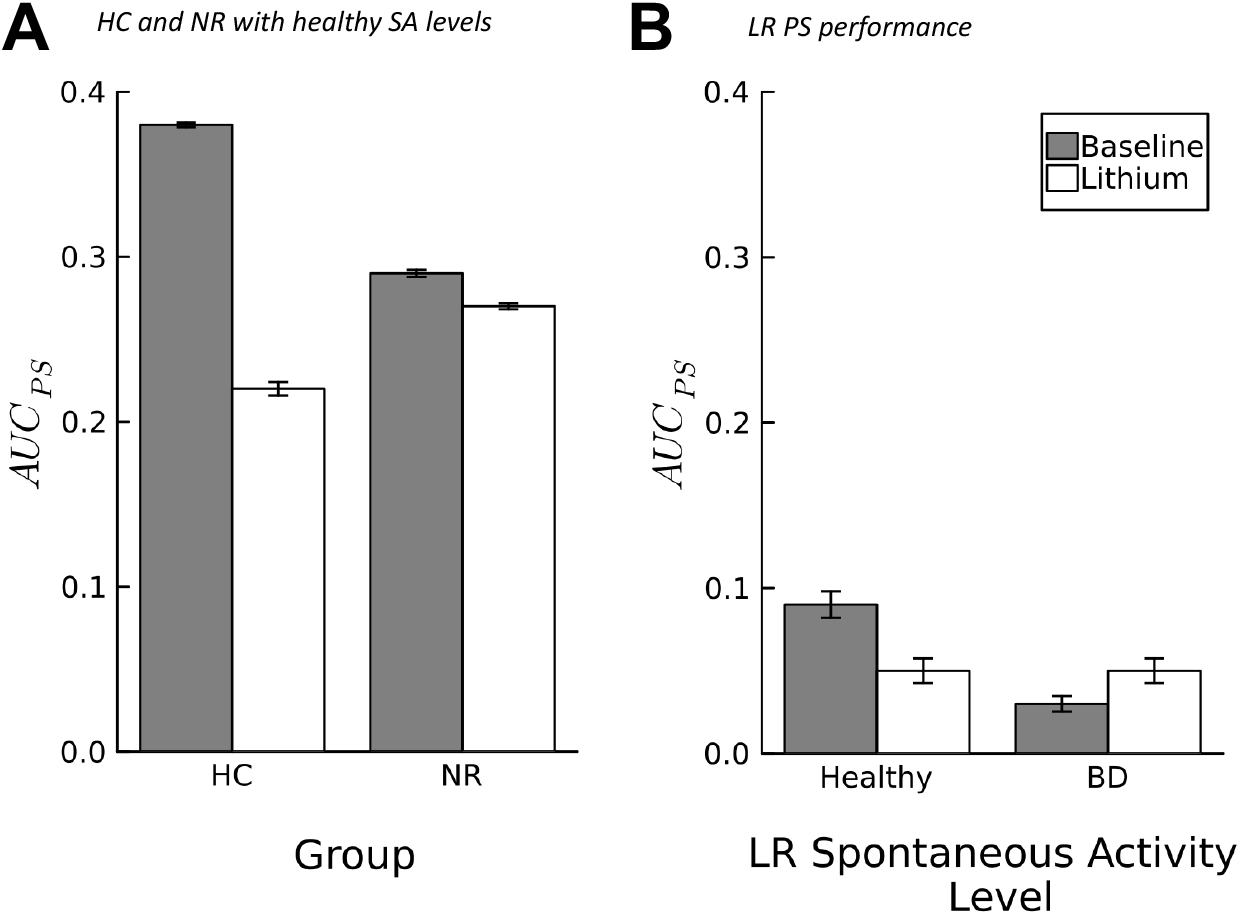
Effects of BD models on pattern separation (PS). A) Healthy levels of spontaneous activity (SA) used for HC and NR models (0.1 Hz [25]), and B) pathological levels of spontaneous activity used for the LR condition (Healthy = 0.1 Hz; BD = 1 Hz, normalised to healthy levels after lithium exposure [25]. Error bars show standard error of the mean, for 14 simulation runs initialised with different random seeds to incorporate variability in network connectivity and granule cells selected for spontaneous activation.

Elevated spontaneous activity levels in LRs without lithium therapy negatively impacted PS (Figure 3B) (LR x Spontaneous Activity; *β* = −0.45, CI: −0.57 – −0.33, p<0.001). Lithium therapy however protected against the deleterious effects of spontaneous activity on PS for LRs (Figure 3B) (LR x Lithium x Spontaneous Activity; *β* = 0.39, CI: 0.22 – 0.56, p<0.001).

### 3.3 Lithium disrupts WTA dynamics in HCs by reducing neuronal sensitivity to negative currents

HC GC population behaviour within the third DG lamella 10 ms post-spontaneous activation is shown in Figure 4 for the GCs directly activated (“AC”) and the remaining GCs within the lamella (“RM”). Without lithium (“CTRL”), GC population activity increased 2 ms post-stimulation, and declined steadily for 3 ms before reaching a steady state of low activity (Figure 4A). The RM population did not show a change in activity levels as well (Figure 4B, “CTRL-RM”). In the lithium-exposed GC model, the directly activated GCs fired more than the CTRL condition post-stimulation, with a similar reduction and stabilisation of activity 3ms post-peak activity (Figure 4A, “LITM-AC”). In the remaining GCs however, there was a substantial and sustained increase in population activity in the lithium-exposed model, despite these GCs not being directly stimulated (Figure 4B, “LITM-RM”). This behaviour was consistent across lamellae, and also for the BD NR models (Supplementary Figure 4). LR models demonstrated similar behaviour, but with a smaller change in the RM activity levels between lithium and control conditions (Supplementary Figure 3). At the cellular level, our lithium-exposed HC GC model demonstrated less sensitivity (i.e., less hyperpolarized response) to negative currents than the control model (Figure 4C). These results align qualitatively with electrophysiological data collected from iPSC GCs *in vitro* (Figure 4D). Lithium increases sensitivity to negative currents in both our LR GC models, and in iPSC GCs *in vitro* (Supplementary Figure 3).

**Figure 4:**
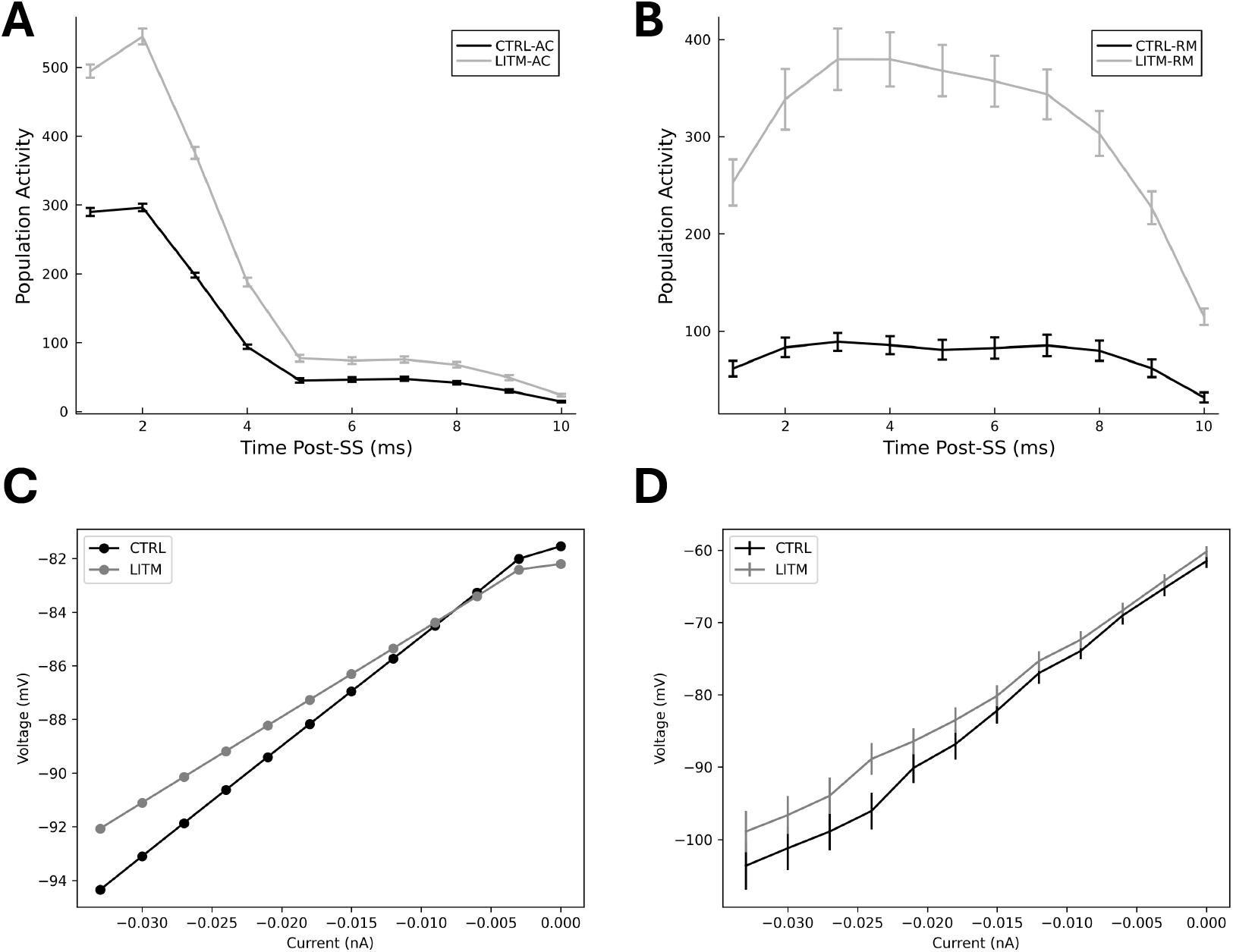
Effects of lithium on healthy control WTA dynamics in DG model (A and B), and cellular response to negative currents in GC model and *in vitro* (C and D respectively). A) GC activity post activation from spontaneous activity generator (“−AC”), and B) activity of remaining GCs not directly stimulated by spontaneous activity (“−RM”) in lamella 3 only. C) HC-CTRL and HC-LITM GC computational model behaviour in response to negative current injection, and D) HC-CTRL and HC-LITM iPSC GC *in vitro* behaviour in response to negative current injection. Error bars show standard error of the mean.

## 4 DISCUSSION

GC hyperexcitability as observed in iPSC GCs derived from BD patients lead disrupted PS relative to HCs in our computational model (Figure 3). Given that PS within the DG is thought to be supported by the intrinsically sparse firing of mature GCs [32, 50, 54, 55], we hypothesised that increased intrinsic excitability of these neurons would lead to PS impairments, which our simulations support. By reducing GC intrinsic excitability, we expected lithium to improve PS in the LR and HC models. Instead, lithium-induced reductions in GC excitability in LRs and HCs impaired PS relative to baseline (i.e., without lithium) models (Figure 3); lithium therefore may not ameliorate PS deficits in LRs by reducing GC intrinsic excitability. Instead, lithium may protect against the loss of PS that would occur with higher spontaneous activity levels in LRs (Figure 3B). Additionally, we identified that lithium not only reduces excitability in HCs, but also sensitivity to negative injected currents (Figure 4C and D). Reducing sensitivity to negative injected currents may prevent effective BC inhibition, leading to inappropriately elevated network activity (Figure 4A and B), which may explain why lithium impairs PS in HCs. Therefore, GC hyperexcitability in BD may lead to PS disruptions, and lithium may prevent these deficits in LRs not by normalising hyperexcitability, but by reducing spontaneous activity levels.

We simulated spontaneous activity by randomly selecting a subset of GCs within the network for stimulation using Poisson spike generators, as the cellular or network mechanism by which this spontaneous activity arises has not yet been identified. Spontaneous activity has been attributable to depolarizing GABA currents [56] and the interplay between sodium and calcium discharges [57] in developing hippocampal circuits, and is thought to tune the development of and promote appropriate synchrony within and between networks [58, 59], raising questions about the implications of elevated spontaneous activity on synaptic plasticity and the functioning of memory systems at the brain network-level in BD. Interestingly, elevated spontaneous activity within the large-scale intrinsic brain networks measured using fMRI may predict diagnosis conversion from major depressive disorder to BD [60], highlighting the importance of understanding spontaneous activity in BD further. Identifying the neural mechanisms of the form of spontaneous activity observed in iPSC GCs, and whether this mechanism also exists *in vivo*, are worthwhile avenues for future investigation. Based on our results, we would predict that treatments that reduce spontaneous activity levels in LRs may preserve PS without the negative effects of lithium therapy.

Indeed, lithium impaired PS for all groups when controlling for spontaneous activity levels (Figure 3A), despite lithium-associated reductions in GC excitability in HCs and LRs, motivating the study of WTA dynamics in our network. In our HC model without lithium, only the directly stimulated GCs fire post-stimulation, with no changes in activity in the remaining GC population (Figure 4A and B), suggesting that BCs are providing sufficient inhibition throughout the network. The elevated and sustained population firing response in our HC-LITM GCs without direct activation (Figure 4B), suggests that BCs are unable to quiet the remaining GCs in the network, or in other words, a deficit in WTA dynamics. Since we did not manipulate the BCs in our network, we hypothesised that this effect may be attributable to change in HC GC sensitivity to inhibition; indeed, our simulations demonstrate that lithium reduces HC GC sensitivity to negative currents (Figure 4C), which agrees with *in vitro* behaviour of iPSC HC GCs (Figure 4D). We present a schematic of this interpretation in Supplementary Figure 5. In summary, although reducing excitability via lithium exposure is theoretically beneficial for neural computations that are reliant on sparse coding such as PS, the effects of lithium on cellular response to inhibition should not be ignored, as it is excitation/inhibition balance within networks that will ultimately promote effective neural computation.

Our cellular and circuit-level results of lithium’s mixed (ie., both beneficial and deleterious) effects for individuals with BD echo clinical discussions of whether lithium is neuroprotective or neurotoxic [61]. There have been reports of lithium-induced cognitive side effects such as memory impairments and a subjective sense of mental “slowness” [62]. Despite these negative effects, lithium is effective at preventing suicide and self-harm in individuals with mood disorders [63] and relapse in individuals with BD [64]. Our results predict that lithium may lead to memory impairments in healthy individuals, motivating a future controlled trial in this group. Finally, our simulations highlight the importance of identifying predictors of lithium response, such that the potential risks to memory systems are mitigated in non-responders, while allowing for excellent responders to benefit from lithium therapy.

One area of promise for identifying predictors may be behavioural tasks that are hypothesised to probe lower-level DG neural computational functioning, such as MD. In a previous study, we hypothesised that PS may underlie MD performance [44], based on two lines of evidence for the DG’s involvement 1) during MD [37, 65], and 2) in PS [33, 34]. Interestingly, a pilot study of the effects of lithium therapy in BD on MD performance demonstrated that lithium therapy significantly improved MD performance in LRs only [39]. Our simulations predict that these MD improvements in LRs may be attributable to lithium-induced reductions in spontaneous activity. We make this statement speculatively, given that 1) a direct empirical demonstration that PS underlies MD has yet to be reported, and 2) the results from our PS simulations must be validated *in vivo*. Further, lithium has been demonstrated to increase hippocampal neurogenesis [66, 67], which has also been shown to improve MD [68–70]. Whether lithium-induced MD improvements are attributable to improvements in PS, increased neurogenesis, or some combination of the two is another avenue for future work, which we discuss further below. Future lines of research addressing these questions will contribute to our understanding of how cellular behaviour impacts neural computation, and how those impacts then translate to behaviour. Forming these mechanistic links across levels of biological hierarchy will allow for the translation of identified cellular-level deficits and drug response to potential mechanistic deficits underlying disease aetiology, observable through patient behaviour.

### 4.1 Strengths and Limitations

#### 4.1.1 Model Fitting procedure

One of our study’s strengths is good model face and predictive validity [71]. Face validity refers to a model’s ability to simulate or capture the behaviour of the system of interest [71]. Our cellular models have face validity because they are directly fit to electrophysiological behaviour of patient- and HC-derived iPSC GCs (Supplementary Figure 1). Predictive validity assesses a model’s ability to predict the effects of interventions and experimental manipulations on the underlying condition [71]. A model with strong predictive validity will be able to generate testable predictions for a set of manipulations in the form of “synthetic” data that can later be compared against real-world data. After model fitting, we assessed our models’ predictive validity by comparing our models’ and iPSC GCs’ membrane response to negative currents (negative IV curve) with and without lithium and found that our models were able to predict lithium-induced 1) reductions in sensitivity to negative currents observed in HCs (Figure 4C and D), and 2) increases in sensitivity to negative currents observed in LRs (Supplementary Figure 3C and D), despite fitting these models to the positive IV and FI data only. Additionally, our simulations are consistent with the effects of lithium on MD performance in individuals with BD [39]. Therefore our fitted GC models exhibit good face and predictive validity, supporting the plausibility of our simulation results.

Although our model fitting procedure was generally successful in producing the iPSC GC behaviour, we had difficulties with fitting the potassium IV curves (Supplementary Figure 1). The potassium channels for all of our computational models were, as a result, more resistant to negative currents than their iPSC counterparts. This issue may be attributable to inaccurate modelling of the potassium channel dynamics and/or a missing additional potassium channel; the baseline GC model used for this study should therefore be re-visited after further genetic analysis, electrophysiology and immunohistochemistry to identify other relevant channel types, their dynamics, and their location on the somatodendritic tree. Additionally, to improve the data available for model fitting purposes, we encourage researchers to follow the electrophysiology protocols outlined by the Allen Brain Institute [72]. Overall, we do not believe that this effect would change the general result of our study as every group/condition was impacted and we were more interested in the relative differences in PS between groups/condition; instead, we believe that this limitation should be addressed in the future, to further improve model face validity.

#### 4.1.2 Improving our Model Design

We made a number of simplifications to our model design that limit biological plausibility. The DG is known to have a subpopulation of adult-born immature GCs that, in contrast to their mature counterparts, are highly intrinsically excitable [73], plastic [74], and are not yet regulated by inhibitory interneurons [75, 76]. Simulations have suggested that these neurons may in fact reduce PS but increase performance on high interference memory tasks [77], which is consistent with studies demonstrating the positive impacts of neurogenesis on discrimination performance [68–70]. As discussed previously, lithium improves MD performance in LRs [39], and also may upregulate neurogenesis in the DG [66, 67]. A natural progression of this model would be to incorporate a sub-population of immature GCs and evaluate PS and performance on a high interference memory task, following previous modellers [77], to test the combined impact of lithium-induced 1) reductions in mature GC intrinsic excitability and spontaneous activity (as we have done here), and 2) upregulated neurogenesis on both PS and MD, to provide us with a more comprehensive and nuanced understanding of lithium’s impacts on DG neural computation.

There have been two iPSC hippocampal neuronal models created to date: the DG GC, and the CA3 pyramidal neuron [25–28]. BD and lithium may also impact the inhibitory interneurons within the hippocampus and DG, of which iPSC neuronal models have not yet been reported. Given that our simulations demonstrated that negative cellular currents modulate network dynamics and PS, motivating future iPSC work on these inhibitory neurons as well. Incorporating detailed models of BD inhibitory interneurons into our network will further improve biological plausibility, and allow for detailed investigation of the complex interplay of abnormal excitatory/inhibitory network dynamics in BD, and subsequent impacts on PS.

Since the time of earlier hippocampal computational models [50, 78], hippocampal research has suggested that the DG performs a number of other computations along with PS, such as contextual binding, novelty detection, temporal tagging, and indexing [79]. We study PS here as a fundamental computation that the DG is ideally suited to perform, but also recognise that future work investigating the impacts of BD on these other computations would be beneficial. PS, although just one of the many computations performed by the DG, seemingly does not conflict with any of the other proposed computations [79], meaning these future results may not contradict the results we have presented here, but rather add to our understanding of DG function in BD. How those other computations then relate to behaviour, and what the clinical implications may be, are yet additional questions.

### 4.2 Conclusions

We presented the first detailed biophysical computational model of GC hyperexcitability and effects of lithium therapy in BD and HCs. We evaluated impacts of the abnormal cellular behaviour on PS using network models, and found that 1) both BD and lithium impair PS in general, 2) lithium may protect against the loss of PS attributable to high spontaneous activity levels in LRs, and 3) lithium reduces sensitivity to negative currents in HCs, impairing GC inhibitory control. Our results are consistent with clinical reports of BD and lithium-associated cognitive slowing and memory impairments, and also with a hypothesised relationship between DG PS and MD. Future work should include a subpopulation of immature GCs to investigate the additional effects of lithium-induced neurogenesis on PS and MD. In conclusion, we presented a first step in translating abnormal iPSC neuronal behaviour derived from BD patients to neural computational deficits; these neural computational deficits may underlie some of the cognitive and memory deficits observed in individuals with BD.

## ACKNOWLEDGEMENTS

This research was supported by Research Nova Scotia (Abraham Nunes), an Ontario Graduate Scholarship (Selena Singh), and the Zuckerman STEM leadership program (Shani Stern). Figure 1 was created using biorender.com.

## AUTHOR CONTRIBUTIONS

Conceptualization and methodology (Selena Singh, Abraham Nunes); Software, formal analysis, and validation (Selena Singh, Abraham Nunes); Resources (Abraham Nunes, Anouar Khayachi); Data Curation (Selena Singh); Writing – Original Draft (Selena Singh, Abraham Nunes); Writing – Review & Editing (All authors); Visualization (Selena Singh, Abraham Nunes); Supervision (Abraham Nunes); Project Administration (Abraham Nunes); Funding Acquisition (Abraham Nunes).

## CODE AVAILABILITY

Code used for our simulations can be found in the following GitHub repository: BDDG-NEURON

