## Supplementary Materials for "The Effects of Bipolar Disorder Granule Cell Hyperexcitability and Lithium Therapy on Pattern Separation in a Computational Model of the Dentate Gyrus"

### 1. Additional Methods

Using an evolutionary algorithm, we fit the parameters of a previously established multicompartmental Hodgkin-Huxley conductance-based model of a mature DG GC [1–3] to both frequency-current (FI), and current-voltage (IV) data from iPSC neurons with and without exposure to lithium ("LITM" and "CTRL", respectively) derived from healthy controls (HC), and LR and NR patients. This procedure generated six models: HC-CTRL, HC-LITM, LR-CTRL, LR-LITM, NR-CTRL and NR-LITM. A mechanism for spontaneous activity was also implemented, following the spontaneous activity levels previously reported in iPSCs[4]. These models therefore behave in a comparable manner to the *in vivo* cell models derived from BD patients. We then integrated each of these computational GC models into a larger DG circuit model, and evaluated the network's PS abilities with each neuronal excitability phenotype.

#### *1.1. Developing GC Models*

We adapted an established multicompartmental Hodgkin-Huxley conductance-based model of the hippocampal dentate GC[1–3,5], implemented in the NEURON simulation environment. This model has two identical dendrites with four compartments each, and a single compartment for the soma. Distributed along the somatodendritic tree are 11 different channels: fast sodium (Na), fast and slow delayed rectifier potassium, A-type potassium, large conductance calcium,voltage-dependent potassium, small conductance calcium-dependent potassium, T-type, N-type, and L-type voltage-gated calcium channels, inward-rectifier potassium and the tonic $GABA_A$ chloride channel. The dynamics of each of these channels are described by sets of differential equations that are parameterized to produce behaviour consistent with what is observed in real-world GCs. Together, these parameters govern the intrinsic excitability and behaviour of these model neurons. To produce our BD and healthy control neuron models, we fit these parameters using numerical optimization to electrophysiological data from iPSC-derived GCs collected from real BD patients and HCs.

##### 1.1.1 Description of iPSC-derived Dentate Gyrus Granule Cell-Like Neurons

iPSC neurons were reprogrammed from lymphocytes and peripheral blood mononuclear cells (PBMC) taken from blood samples from consenting participants. Detailed methodology describing iPSC differentiation and cell culture protocols for the iPSC neurons used to inform our modelling have been previously described[6]. We therefore provide a brief overview here.

Blood samples were collected from 8 BD patients (4 LRs, and 4 NRs), and 5 HC participants. After lymphocyte and PBMC isolation, and iPSC differentiation as described previously[6,7], about half of the neurons per group were exposed to therapeutic levels of lithium (~1.5 mM), for 7 days. Therefore, there were the following number of iPSC GC-like neurons per group used for whole-cell patch-clamp recordings: LR $\{n_{Li} = 49 , n_{CTRL} = 55\}$; NR $\{ n_{Li} = 41 , n_{CTRL} = 45\}$; HC $\{n_{Li} = 42, n_{CTRL} = 40\}$.

Sodium and potassium current-voltage (IV), and frequency-current (FI) relationships were acquired in voltage-clamp and current-clamp modes respectively. Sodium channel currents were reported as inward peak currents, and potassium channel currents as outward currents, and voltages were stepped up by 10mV from -70mV to 20mV. Frequency-current relationships were acquired following a current-clamp protocol, where current was injected for 1s from

0pA to 33pA in 3pA steps, and the number of resulting action potentials were recorded. Current-voltage relationships for negative current values were acquired in current-clamp mode, where current was injected for 1s from -33pA to 0pA in 3pA steps, and the membrane potential was recorded. These negative current-voltage curves were used after our parameter optimization procedure for further model validation.

##### *1.1.2 Numerical Optimization-Based Fitting of Computational Models to Cellular Data*

To develop biophysically realistic models of HC LR and NR GCs, a custom evolutionary algorithm was built using the *inspyred* (v. 1.0) and *NetPyNe* (v. 1.0.0.2) Python packages to facilitate parameter optimization, such that the resulting models produce FI and IV curves that align with average FI and IV curves of iPSC GCs. The iPSC neurons were excitable for current and voltage injection values that were one order of magnitude smaller than the values used in the original GC model [1–3]. To accommodate for this large difference in excitability, sodium and fast/slow potassium channel activation and inactivation thresholds were also optimised, along with the 11 channel conductances outlined earlier, for all neuronal compartments.

Evolutionary algorithms perform parameter optimization by iteratively mutating, then evaluating the "fitness" of a parameter set[8]. Here, we computed fitness as the summed mean squared error (MSE) between the simulated FI and IV curves and FI and IV curves from iPSC data; our goal was to find parameter sets that *minimised* the summed MSE. We deemed the model fits to be satisfactory if each simulated data point fell within the empirical standard error of the mean. 350 evaluations were made to ensure stable convergence to a minimum MSE value.

##### *1.1.3 Granule Cell-Like Neuron Models for Lithium Nonresponders*

Experimental data failed to show a statistically significant effect of lithium exposure on FI and IV curves for the NR iPSC GCs, meaning these two curves were statistically identical. Therefore, to produce a NR-LITM model, we began with the fitted NR-CTRL model and modified the parameters by increasing or decreasing their values by a random value less than 2% of the original parameter's value to introduce some noise. This approach yielded two models with slight differences that have comparable parameter values and biophysical behaviour, which we believe are good candidates for simulating NR-CTRL and NR-LITM conditions.

##### *1.1.4 Simulation of Spontaneous Activity*

To simulate spontaneous activity, randomly selected GCs within the network were equipped with Poisson spike generators, synapsed onto GC somata, that randomly produced spikes at the following rates during the simulation: HC and NRs = 0.25 Hz; LRs = 1Hz, following previous experimental reports of spontaneous activity levels [9]. The effect of lithium on spontaneous activity was captured by setting the LR spontaneous activity level back to HC levels of 0.25 Hz[9].

#### ***1.2. Biophysical Network Model of the Dentate Gyrus***

We employed a previously established conductance-based biophysical model of the DG [2,3,5] to study the impacts of BD GC hyperexcitability on PS. This network was implemented in the NEURON simulation environment (v. 8.0)[10], and used the original model's geometric and topological features. Our model included 500 glutamatergic GCs (as described earlier), 6 GABAergic basket cells (BCs), 15 glutamatergic mossy cells (MCs), 6 GABAergic hilar perforant path cells (HIPP), and 100 excitatory entorhinal perforant path (PP) cells . As with the GCs, the other cells in the network (BCs, MCs, HIPP) were modelled as multicompartmental Hodgkin-Huxley style neurons with a soma and varying numbers of dendrites: BCs had 4 dendrites with 4 compartments each; MCs had 4 dendrites with 4 compartments each; and HIPP cells had 4 dendrites with 3 compartments each. Ion channel density varied for each cell and cellular compartment, and the same ion channels described earlier are also included in these models with different conductance values (see Section 3 for additional detail). Connectivity in our model was topographically distributed, and synaptic strengths were randomly initialised [2,3]. Excitatory glutamatergic AMPA synapses and inhibitory GABAergic synapses were modelled using double-exponential functions.

PP cells were modelled as point processes that stimulated GCs and BCs. All biophysical properties were kept the same as in the original model [2,3,5]. Parameter values for the connectivity, cellular biophysics, and synaptic double-exponential functions can be found in section 3, and also described in our previous study that employed this model[5].

*1.2.1 Spatiotemporal PS task*

The present study adopted the same spatiotemporal PS task first presented by Myers and Scharfman in 2009, and later adapted by Yim *et al.*[3] This protocol involved simulating 24 partially overlapping patterns of PP inputs over a 200 ms window. A single "pattern" consisted of spike trains from 6 of the 100 PP cells to align with activity levels within the entorhinal cortex [11,12]. The $i^{th}$ input pattern therefore consisted of spike trains generated by PP cells $\{i, i+1, ..., i+5\}$.

To calculate Pearson correlations of PP input patterns and GC output patterns, PP and GC spike trains first needed to be flattened into single time series representations, which was done by convolving spike trains with a triangular kernel. Pearson correlations between these time series representations were then calculated for PS analysis, as we have done in our previous study[5]. For example, let the correlation between PP patterns $i$ and $j$ be denoted $x = Corr(PP_i, PP_j)$, and the resulting correlation between GC population activity driven by patterns $i$ and $j$, respectively, was $y = Corr(GC_i, GC_j)$. Each of the 156 $(x, y)$ pairs for the set of 24 entorhinal cortex patterns was plotted and fit with a *pattern separation curve* defined as $y = a + (1-a)x^b$, where $0 \leq a \leq 1$ and $0 \leq b$ are parameters describing the shape of this curve. After fitting this equation using numerical least-squares optimization, we computed a summary PS index defined as the area between the leading diagonal and the PS curve:

$$AUC_{PS} = -\frac{2ab - b + 1}{2\,(1+b)}$$

Higher values of $AUC_{PS}$ indicate stronger PS by the DG network. PS analyses were conducted using custom Julia scripts (v. 1.8.5).

### 2. Additional Results

#### *2.1. Results from parameter optimization procedure*

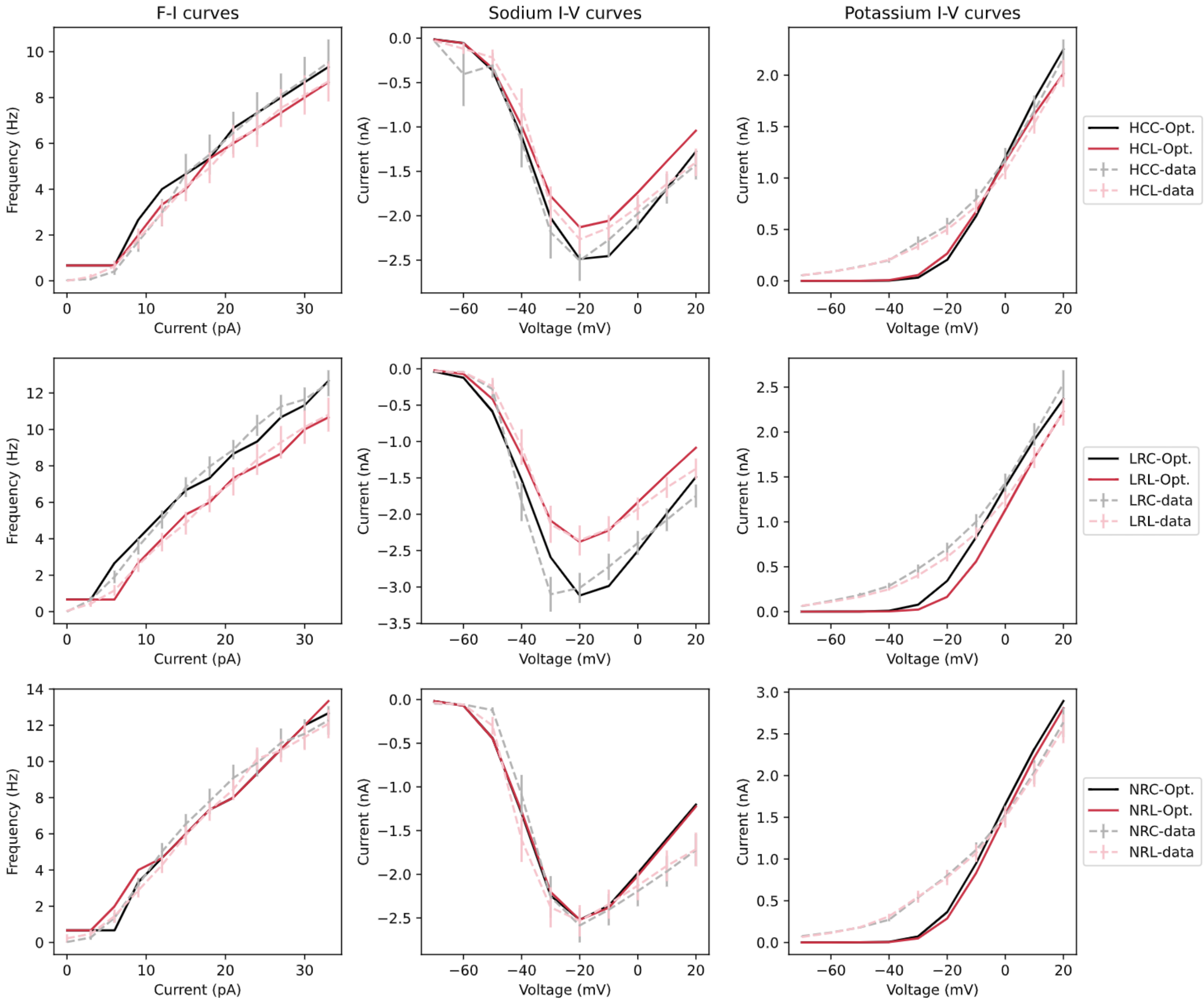

*Figure 1: Parameter optimization produces simulated (solid black = baseline, and red = with lithium) FI and sodium/potassium IV relationships that align with iPSC data (dashed grey = baseline, pink = with lithium). HCC: Healthy controls, control condition; HCL: Healthy controls, lithium condition; LRC: LRs, control condition; LRL: LRs, lithium condition; NRC: NRs, control condition; NRL: NRs, lithium condition. "Opt." refers to the simulated curve after optimization, and "data" refers to raw data from iPSCs.*

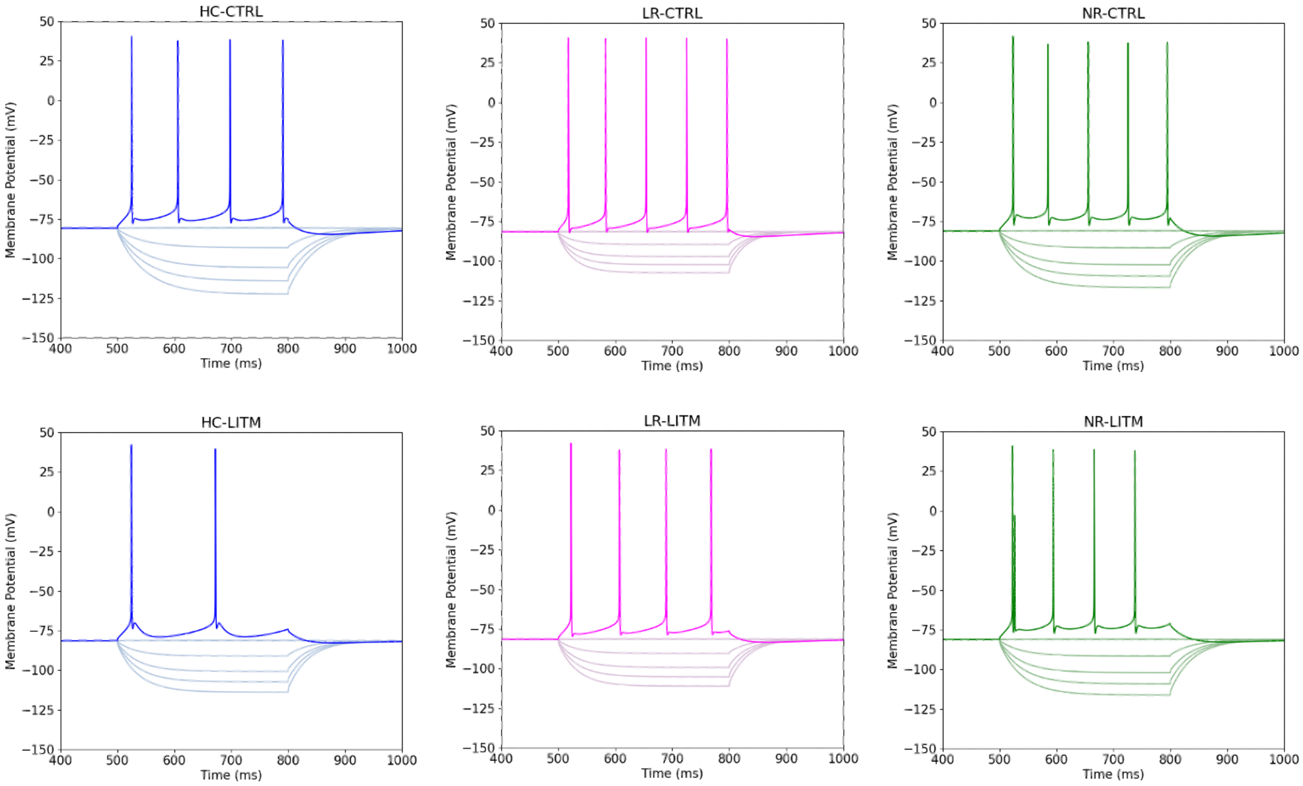

*Figure 2: HC, BD LR, NR, membrane currents, with and without lithium, from simulated GCs. Current injection values were as follows, in pA: -100, -80, -60, -30, 0, 50. Currents were injected and membrane potential was recorded from the soma.*

*2.2. Statistical analysis*

*Table 1: Regression table for linear mixed effects model.*

| | zAUC | | |
|---|---|---|---|
| *Predictors* | *Estimates* | *CI* | *p* |
| (Intercept) | 1.37 | 1.29 – 1.44 | **<0.001** |
| Group [LR] | -2.37 | -2.46 – -2.29 | **<0.001** |
| Group [NR] | -0.75 | -0.83 – -0.66 | **<0.001** |
| Lithium | -1.32 | -1.41 – -1.24 | **<0.001** |
| SpontActiv | 0.00 | -0.09 – 0.09 | 1.000 |
| Group [LR] × Lithium | 1.00 | 0.88 – 1.13 | **<0.001** |
| Group [NR] × Lithium | 1.22 | 1.09 – 1.34 | **<0.001** |
| Group [LR] × SpontActiv | -0.45 | -0.57 – -0.33 | **<0.001** |
| Group [NR] × SpontActiv | -0.00 | -0.12 – 0.12 | 1.000 |
| Lithium × SpontActiv | 0.06 | -0.06 – 0.18 | 0.326 |
| (Group [LR] × Lithium) × SpontActiv | 0.39 | 0.22 – 0.56 | **<0.001** |
| (Group [NR] × Lithium) × SpontActiv | -0.06 | -0.23 – 0.11 | 0.487 |
| **Random Effects** | | | |
| $\sigma^2$ | 0.01 | | |

| | |
|---|---|
| $\tau_{00\ ID}$ | 0.01 |
| ICC | 0.32 |
| $N_{ID}$ | 14 |
| Observations | 168 |
| Marginal $R^2$ / Conditional $R^2$ | 0.980 / 0.987 |

*2.2. WTA dynamics and negative current injection protocol for lithium responders*

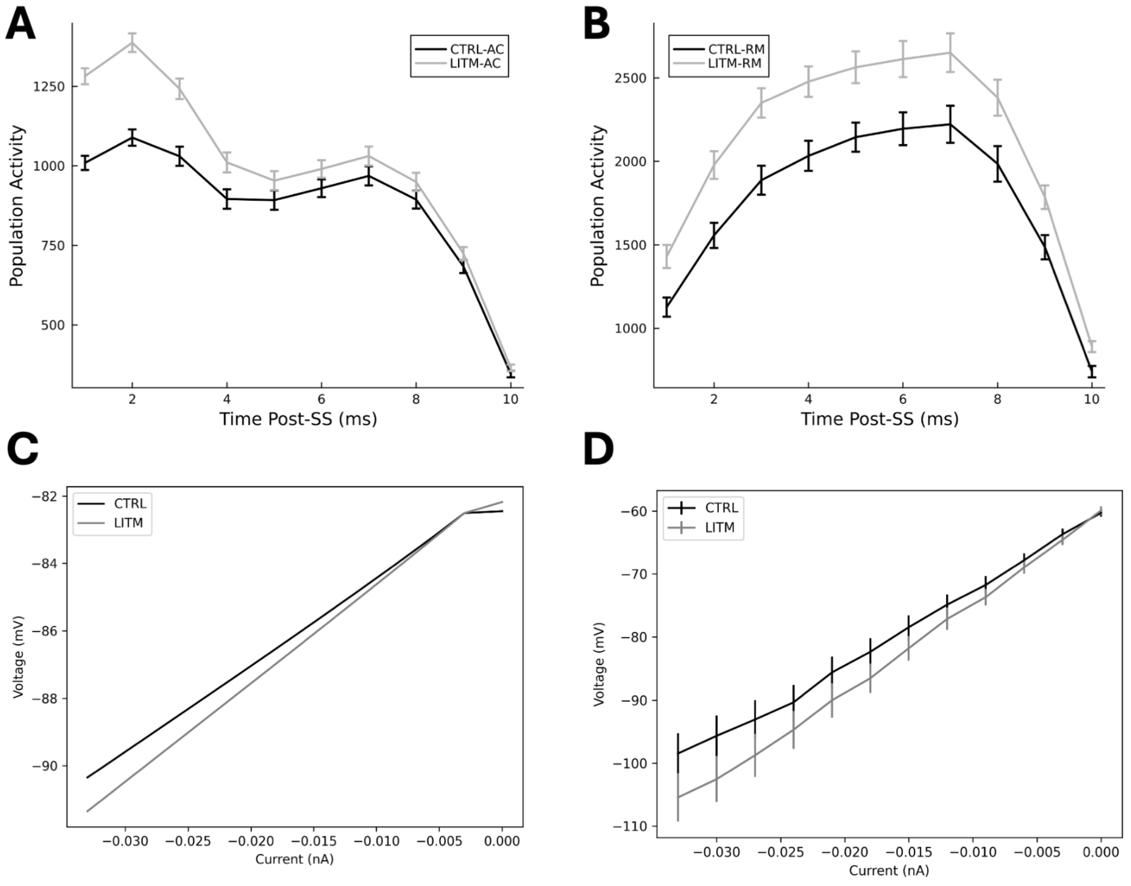

*Figure 3: Effects of lithium on lithium responder WTA dynamics in DG model (A and B), and cellular response to negative currents in GC model and in-vitro (C and D respectively). A) GC activity post activation from spontaneous activity generator ("-AC"), and B) activity of remaining GCs not directly stimulated by spontaneous activity ("-RM"). C) LR-CTRL and LR-LITM GC computational model behaviour in response to negative current injection, and D) LR-CTRL and LR-LITM iPSC GC in-vitro behaviour in response to negative current injection. Error bars show standard error of the mean.*

*2.3. WTA dynamics and negative current injection protocol for lithium nonresponders*

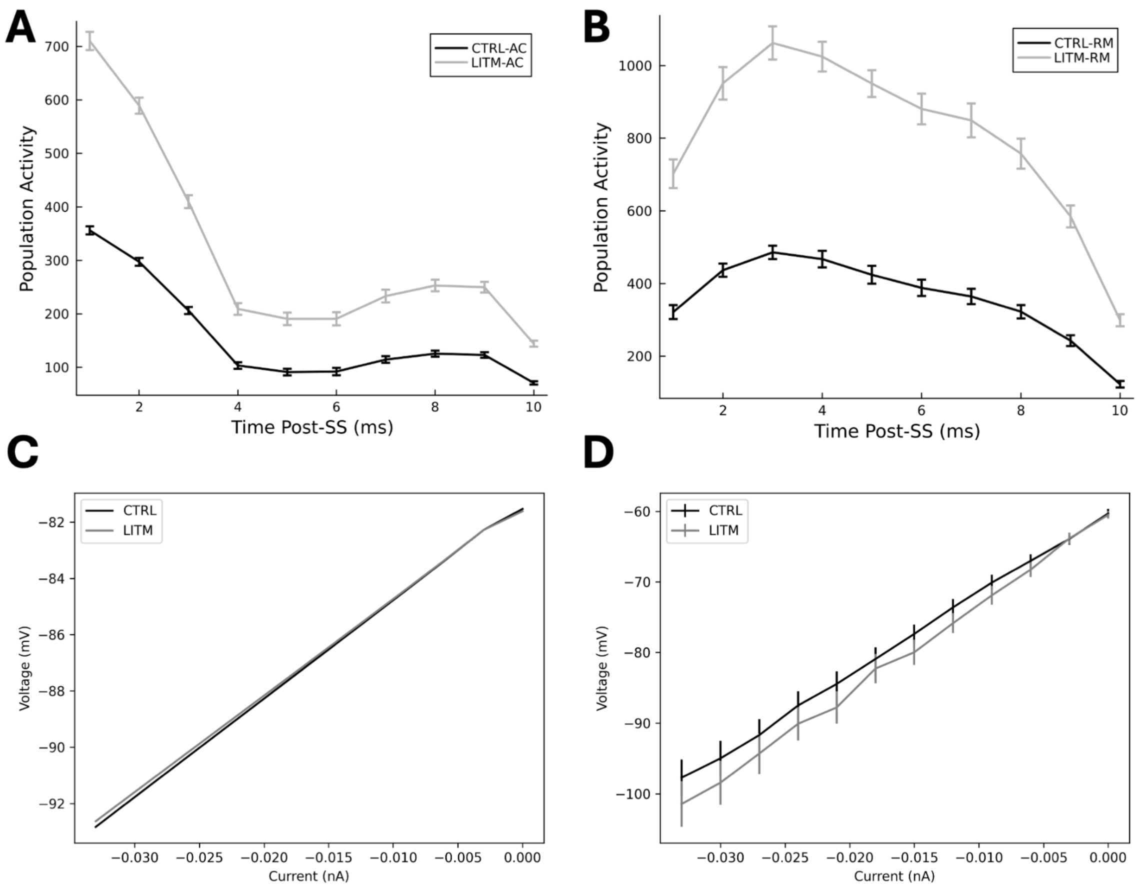

*Figure 4: Effects of lithium on lithium non-responder WTA dynamics in DG model (A and B), and cellular response to negative currents in GC model and in-vitro (C and D respectively). A) GC activity post activation from spontaneous activity generator ("-AC"), and B) activity of remaining GCs not directly stimulated by spontaneous activity ("-RM"). C) NR-CTRL and NR-LITM GC computational model behaviour in response to negative current injection, and D) NR-CTRL and NR-LITM iPSC GC in-vitro behaviour in response to negative current injection. Error bars show standard error of the mean.*

**3. Parameter Tables for Biophysical Model**

We provide a summary of the relative differences in sodium and potassium maximum conductance values between groups below. Percentage changes in sodium and potassium conductances in BD models relative to HC models are shown in Table 1 below, and in the HC and BD-LRs with and without lithium in Table 2 below.

Table 1: Changes in Na and K maximum conductances relative to HC in BD models, without lithium:

| Comparison | Soma | | Dendrites | |
|---|---|---|---|---|
| HC vs. LR | *Na* | 0% | *Na* | 108.79% |
| | *K* | 16.62% | *K* | 70.02% |
| HC vs. NR | *Na* | 10.32% | *Na* | 41.29% |
| | *K* | 10.47% | *K* | 63.82% |
| LR vs NR | *Na* | -9.36% | *Na* | 119.56% |
| | *K* | 7.96% | *K* | 84.91% |

Table 2: Changes in Na and K maximum conductances after lithium administration in HC and BD-LR models:

| Group | Soma | | Dendrites | |
|---|---|---|---|---|
| HC | *Na* | 12.90% | *Na* | 91.54% |
| | *K* | -6.96% | *K* | 51.64% |
| LR | *Na* | 12.90% | *Na* | 26.02% |
| | *K* | -7.57% | *K* | 52.76% |

*3.1. Parameters for HC, LR, and NR GC models, with and without lithium, after model fitting*

*Table 3: Conductance and biophysical parameter values for all model neurons, for each compartment. HC: healthy control; LR: lithium responder; NR: lithium nonresponder; CTRL: control condition; LITM: lithium exposed condition.*

| Cellular Section | Parameter | Baseline Value (Yim et al., 2015) | HC - CTRL | HC - LITM | LR - CTRL | LR - LITM | NR - CTRL | NR - LITM |
|---|---|---|---|---|---|---|---|---|
| soma | gkbar | 0.0006 | 6E-05 | 0.0012 | 0.0009053947902481570 | 0.00010788516644276900 | 0.0010226409107385500 | 0.0003660975416788850 |
| soma | gnatbar | 0.12 | 0.27 | 0.3048255894843410 | 0.27 | 0.3048255894843410 | 0.2978835265729490 | 0.30382538214974500 |
| soma | vshiftma | 43.0 | 66.9197734992995 | 67.7915876735032 | 70.7380175192901 | 68.32860143407120 | 66.9197734992995 | 67.29851063919570 |
| soma | vshiftmb | 15.0 | 20.0733027547385 | 20.814837386665500 | 19.8 | 22.68359220456880 | 21.27948167559500 | 23.470953358463600 |
| soma | vshiftha | 65.0 | 118.06045177335300 | 129.0173357056420 | 117.23032143397900 | 131.61636329083800 | 122.15522840474800 | 118.87882140094200 |
| soma | vshifthb | 12.5 | 20.942582083983100 | 19.77649734464790 | 19.227705061051100 | 18.0 | 18.237340642992300 | 22.726632076322700 |
| soma | vshiftnfa | 18.0 | 32.70452482541600 | 35.465023068663600 | 36.300000000000000 | 30.944730312092700 | 33.525987070921000 | 33.30911097464430 |
| soma | vshiftnfb | 43.0 | 70.83253515067660 | 76.0102085023672 | 76.0102085023672 | 76.0102085023672 | 83.13767945881000 | 76.56629644409440 |
| soma | vshiftnsa | 30.0 | 39.00392171045480 | 43.51613735432970 | 44.419782716630500 | 41.644727839531800 | 41.962683474184200 | 43.38534894819660 |
| soma | vshiftnsb | 55.0 | 100.52203817524900 | 97.79072832014910 | 96.44920629762610 | 97.86371169177150 | 109.05837400839700 | 107.92862157059400 |
| soma | gkfbar | 0.016 | 0.023981290697889000 | 0.020641914259992600 | 0.0239812906978890000 | 0.0239812906978890000 | 0.029003294615369500 | 0.0295926683090268000 |
| soma | gksbar | 0.006 | 0.0009000000000000000 | 0.0009000000000000000 | 0.0011991879551190100 | 0.0010177168001500000 | 0.0009000000000000000 | 0.0008982952385591360 |
| soma | gl | 1.44E-05 | 1.84584649593557E-05 | 1.44E-05 | 2.69437340439673E-05 | 1.44E-05 | 2.88E-05 | 1.48295202123238E-05 |
| soma | glcabar | 0.005 | 0.002899776815334780 | 0.006072037456519690 | 0.0005 | 0.0005 | 0.0005 | 0.0025535189223371400 |
| soma | gncabar | 0.002 | 0.0002 | 0.0020484879674828300 | 0.0012769067146698600 | 0.0002 | 0.0002 | 0.00019822407216141600 |
| soma | gskbar | 0.001 | 0.0005759620485886370 | 0.0005759620485886370 | 0.0001 | 0.0010316073872960700 | 0.0014651489032659300 | 0.0005111892553052610 |

| | | | | | | | | |
|---|---|---|---|---|---|---|---|---|
| soma | gcatbar | 3.7E-05 | 1.85E-06 | 1.37396973464378E-05 | 1.85E-06 | 1.85E-06 | 1.85E-06 | 1.93161941306084E-06 |
| gcdend1_0 | gkbar | 0.0006 | 0.0011388 | 0.00015396934171632200 | 0.0013392 | 0.000111000000000000000 | 0.0004669114281027840 | 0.00043569687585868800 |
| gcdend1_0 | gnatbar | 0.018 | 0.006585161639596600 | 0.0018 | 0.0054000000000000000 | 0.0044230950761171400 | 0.0045 | 0.0058756670090355400 |
| gcdend1_0 | vshiftma | 43.0 | 24.159513535551500 | 8.8335926839605 | 27.29424126470690 | 21.43191595408170 | 73.40115270584590 | 72.00908352837670 |
| gcdend1_0 | vshiftmb | 15.0 | 5.130917449198990 | 19.250171535316400 | 5.527870229166840 | 20.532250384002500 | 11.981794722934300 | 10.058662872976300 |
| gcdend1_0 | vshiftha | 65.0 | 95.30762369177960 | 94.26226772828050 | 67.0506246711742 | 54.18556939012500 | 57.121821371553300 | 105.82526035660300 |
| gcdend1_0 | vshifthb | 12.5 | 10.528032531247600 | 12.096146624908300 | 13.28422638916720 | 15.390824100064500 | 25.627217427564000 | 16.077005136300600 |
| gcdend1_0 | vshiftnfa | 18.0 | 28.868924308694200 | 24.576664932784200 | 33.77419384505460 | 5.710416464626170 | 31.669906114570900 | 26.6930361914663 |
| gcdend1_0 | vshiftnfb | 43.0 | 71.30868858993190 | 40.455754618082200 | 20.47255762760280 | 75.71966597348600 | 83.93425688905380 | 20.320443221665000 |
| gcdend1_0 | vshiftnsa | 30.0 | 13.827615914797500 | 31.653652485856800 | 42.78767652707100 | 35.50770698285330 | 13.27485916607510 | 36.397948927163600 |
| gcdend1_0 | vshiftnsb | 55.0 | 38.40857344961370 | 18.69555630109860 | 60.19766316264460 | 51.08049947727820 | 56.28786497488660 | 55.5066956985511 |
| gcdend1_0 | gkfbar | 0.004 | 0.0030517109412500600 | 0.006893185159071110 | 0.0074369239764473900 | 0.0010499438330151800 | 0.0013347321022055300 | 0.0060453088497373000 |
| gcdend1_0 | gksbar | 0.006 | 0.005185245531565050 | 0.002593414024361270 | 0.0133920000000000000 | 0.00111 | 0.007270158534437920 | 0.0072228904944001900 |
| gcdend1_0 | gl | 1.44E-05 | 2.73312E-05 | 2.808E-05 | 3.21408E-05 | 2.73312E-05 | 2.9350393637413E-05 | 2.4438552923756E-05 |
| gcdend1_0 | glcabar | 0.0075 | 0.007966962171984310 | 0.0146250000000000000 | 0.00225 | 0.0022979109034021300 | 0.001875 | 0.0023877302702145300 |
| gcdend1_0 | gncabar | 0.003 | 0.0009 | 0.003164489854829350 | 0.0009 | 0.000555 | 0.0036144898548293400 | 0.0010788454698276800 |
| gcdend1_0 | gskbar | 0.0004 | 0.00012 | 0.00023276430969005000 | 0.00012 | 7.4E-05 | 0.0001 | 0.0001234648752524550 |
| gcdend1_0 | gcatbar | 7.5E-05 | 0.0001369229136409590 | 7.5E-06 | 0.000114452672393350000 | 8.16549578982843E-05 | 0.00013044761099805700 | 2.3664058927393E-05 |
| gcdend1_1 | gkbar | 0.001 | 0.0009446537767857340 | 0.00195 | 0.0003 | 0.000185 | 0.0021000000000000000 | 0.0019006128979532300 |
| gcdend1_1 | gnatbar | 0.013 | 0.0039 | 0.0013 | 0.0234540683064015000 | 0.002405 | 0.00325 | 0.0241737589121375000 |

| | | | | | | | | |
|---|---|---|---|---|---|---|---|---|
| gcdend1_1 | vshiftma | 43.0 | 36.112243133627700 | 36.613775509579500 | 52.278410397882000 | 52.11818752582000 | 41.87070851163090 | 35.739286450342000 |
| gcdend1_1 | vshiftmb | 15.0 | 24.655134533085300 | 13.82089074040400 | 9.165188985074990 | 24.38059790686800 | 13.434857297344100 | 8.643763774347960 |
| gcdend1_1 | vshiftha | 65.0 | 121.34626450273200 | 124.09470709870800 | 142.0948952915690 | 103.86736398057 | 115.43760850207500 | 116.92605508197800 |
| gcdend1_1 | vshifthb | 12.5 | 19.521435597348700 | 17.81115188729120 | 12.049185986835100 | 9.626287516332530 | 11.027290870471500 | 5.985162128333980 |
| gcdend1_1 | vshiftnfa | 18.0 | 25.20814805905520 | 4.308808187778060 | 40.176 | 5.653020770629090 | 29.83788436831300 | 33.04698698782420 |
| gcdend1_1 | vshiftnfb | 43.0 | 16.025645267730700 | 80.60460188253270 | 17.176401155174000 | 59.486489550863400 | 38.569357411019700 | 44.256614222258800 |
| gcdend1_1 | vshiftnsa | 30.0 | 52.14250357840310 | 12.03859361197070 | 60.078022548235300 | 51.85703960706900 | 57.51052146706230 | 50.935451524790500 |
| gcdend1_1 | vshiftnsb | 55.0 | 68.58907801718650 | 5.5 | 69.33591626979850 | 76.45373690487040 | 38.49769452572980 | 61.792597223820400 |
| gcdend1_1 | gkfbar | 0.004 | 0.003278706286681320 | 0.001791209575976130 | 0.002278311790858880 | 0.007592 | 0.008400000000000000 | 0.003313350142465580 0 |
| gcdend1_1 | gksbar | 0.006 | 0.010146078173457800 | 0.0117 | 0.009665999167902250 | 0.011388 | 0.0126 | 0.008212484290698330 |
| gcdend1_1 | gl | 2.268E-05 | 4.304664E-05 | 4.08563045346615E-05 | 4.78598897208902E-05 | 4.304664E-05 | 5.67E-06 | 4.20300062273966E-05 |
| gcdend1_1 | glcabar | 0.0075 | 0.014235 | 0.014625000000000000 | 0.00225 | 0.001680199140903090 0 | 0.001875 | 0.002686165278841090 0 |
| gcdend1_1 | gncabar | 0.001 | 0.0018828710352034900 | 0.0001 | 0.0003 | 0.001428787461090523 0 | 0.00025 | 0.000997150921925724 0 |
| gcdend1_1 | gskbar | 0.0002 | 0.0003714695426649100 | 2E-05 | 0.00044640000000000000 | 3.7E-05 | 8.75550811684428E-05 | 9.96535231981931E-05 |
| gcdend1_1 | gcatbar | 0.00025 | 0.00043685939897947000 | 0.0004511437599711120 | 0.000558 | 0.000390211607719104 0 | 0.000433970504541940 00 | 0.00045442618393490600 |
| gcdend1_2 | gkbar | 0.0024 | 0.0024380529003697000 | 0.002254446525098730 | 0.000720000000000000 00 | 0.000714659525222440 0 | 0.005040000000000000 | 0.001554386208835950 0 |
| gcdend1_2 | gnatbar | 0.008 | 0.0024 | 0.012335056518284300 | 0.003643370571547990 | 0.010415754606233700 | 0.009924161304063 | 0.01092949038241920 |
| gcdend1_2 | vshiftma | 43.0 | 34.892232105469300 | 13.06300945059780 | 62.63047856843870 | 20.57885443999990 | 24.383468017513000 | 53.42393602450570 |
| gcdend1_2 | vshiftmb | 15.0 | 18.18955414218300 | 13.158083471076300 | 16.1649466206203 | 7.280398266819820 | 17.845352829945900 | 26.316020776834700 |
| gcdend1_2 | vshiftha | 65.0 | 20.315483923757100 | 115.92274169394700 | 78.33034025457030 | 100.52091020515100 | 128.04265312097900 | 71.56344028728050 |

| | | | | | | | | |
|---|---|---|---|---|---|---|---|---|
| gcdend1_2 | vshifthb | 12.5 | 10.200506306287800 | 10.457812728809100 | 12.152305533506000 | 9.227216710056980 | 5.646962687156770 | 5.104871273921290 |
| gcdend1_2 | vshiftnfa | 18.0 | 22.75832918678170 | 22.48403510289840 | 27.644354998764200 | 19.112814932178300 | 7.486947371022530 | 23.30935492611520 |
| gcdend1_2 | vshiftnfb | 43.0 | 14.655422764890300 | 39.981697919205900 | 15.022325895975000 | 40.99432353275660 | 89.28589602641860 | 81.83265112336160 |
| gcdend1_2 | vshiftnsa | 30.0 | 48.357390613300100 | 28.663616096364700 | 36.329813207241800 | 47.844432909109800 | 14.705168550198200 | 19.762014624596900 |
| gcdend1_2 | vshiftnsb | 55.0 | 86.64144691534450 | 100.71304777942400 | 53.898751811717900 | 65.1406415681786 | 54.00035166029620 | 83.33946866528910 |
| gcdend1_2 | gkfbar | 0.001 | 0.001898 | 0.000782102632881809 | 0.0011030187917170400 | 0.0018564710206225400 | 0.0020551496720091600 | 0.0011736442106936800 |
| gcdend1_2 | gksbar | 0.006 | 0.006851227062029810 | 0.002926463742229500 | 0.0079069904154202800 | 0.00652473839628102 | 0.010805424873646900 | 0.0058808339164698000 |
| gcdend1_2 | gl | 2.268E-05 | 9.2125709528675E-06 | 2.427558372151E-05 | 5.062176E-05 | 3.98795996334507E-05 | 1.3305052037089E-05 | 1.32762743797171E-05 |
| gcdend1_2 | glcabar | 0.0005 | 0.000554979371694051 | 0.0005935457843622790 | 0.0006396246220981890 | 0.0007214102561293200 | 0.0006685457843622790 | 0.00016453535694872600 |
| gcdend1_2 | gncabar | 0.001 | 0.0003 | 0.0013696485661433800 | 0.0003 | 0.0002375905491156150 | 0.0021000000000000000 | 0.0014033105293291900 |
| gcdend1_2 | gskbar | 0.0 | 0.0 | 0.0 | 0.0 | 0.0 | 0.0 | 0.0 |
| gcdend1_2 | gcatbar | 0.0005 | 0.00045753601563215600 | 0.000975 | 0.00052181450700959 | 0.0001422206333873960 | 0.0008054324575045870 | 0.0004553956602161600 |
| gcdend1_3 | gkbar | 0.0024 | 0.0045552 | 0.0046800000000000000 | 0.0053568 | 0.0045552 | 0.0006 | 0.00416445454329038 |
| gcdend1_3 | gnatbar | 0.0 | 0.0 | 0.0 | 0.0 | 0.0 | 0.0 | 0.0 |
| gcdend1_3 | vshiftma | 43.0 | 71.82070754626790 | 24.55654634386180 | 46.801086453664700 | 63.720561975428300 | 60.140749759953800 | 62.09792868319230 |
| gcdend1_3 | vshiftmb | 15.0 | 23.82096357263450 | 20.75843648652360 | 28.067116081644400 | 23.746590604742300 | 26.398828148729200 | 23.332657237378600 |
| gcdend1_3 | vshiftha | 65.0 | 25.966817283953900 | 92.17576694194550 | 27.318454939048200 | 18.957201506516300 | 88.01416506469670 | 82.6490279771471 |
| gcdend1_3 | vshifthb | 12.5 | 12.77094555892020 | 15.386113981477200 | 15.466097103804400 | 12.654954648962900 | 13.3535120292912 | 11.902673776089300 |
| gcdend1_3 | vshiftnfa | 18.0 | 9.062255863198690 | 24.597714363821500 | 21.971841778118200 | 7.674914562947120 | 28.712428479598300 | 10.17372302817700 |
| gcdend1_3 | vshiftnfb | 43.0 | 33.182284275349500 | 10.054466563705600 | 31.732251089655700 | 34.2740120449535 | 18.878005128625100 | 20.529083779324700 |

| | | | | | | | | |
|---|---|---|---|---|---|---|---|---|
| gcdend1_3 | vshiftnsa | 30.0 | 28.650864667823200 | 45.04701766913720 | 52.56354362251170 | 26.61503828284170 | 53.63934757923690 | 27.767585507950400 |
| gcdend1_3 | vshiftnsb | 55.0 | 40.538546431288900 | 33.52917452469620 | 22.40030389623540 | 37.98813856513700 | 42.078198698264700 | 65.82531822034420 |
| gcdend1_3 | gkfbar | 0.001 | 0.0003 | 0.0001 | 0.0003694241661871250 | 0.0015016704865968900 | 0.0011909102406158700 | 0.00036926305629303100 |
| gcdend1_3 | gksbar | 0.008 | 0.0024 | 0.0156 | 0.00952359150433421 | 0.0086942728932597100 | 0.002 | 0.0062060166277206900 |
| gcdend1_3 | gl | 2.268E-05 | 1.87711665133128E-05 | 4.4226E-05 | 5.062176E-05 | 4.304664E-05 | 4.33555478259339E-05 | 4.33833350061444E-05 |
| gcdend1_3 | glcabar | 0.0 | 0.0 | 0.0 | 0.0 | 0.0 | 0.0 | 0.0 |
| gcdend1_3 | gncabar | 0.001 | 0.0009738924082332250 | 0.00195 | 0.002232 | 0.0009073890458720360 | 0.0021000000000000000 | 0.0003306300959241470 |
| gcdend1_3 | gskbar | 0.0 | 0.0 | 0.0 | 0.0 | 0.0 | 0.0 | 0.0 |
| gcdend1_3 | gcatbar | 0.001 | 0.001898 | 0.00195 | 0.002232 | 0.00020630341415461600 | 0.0005300414789396060 | 0.0018542532259645000 |

*3.2. Parameters for BC, HIPP and MC models*

*Table 4: Basket cell biophysics. † indicates manual adjustment for excitability, to ensure compatibility with iPSC fitted models.*

| **Segment(s)** | **Param** | **Value** |
|---|---|---|
| All | $\tau^{\text{ccanl}}_{\text{Ca}}$ | 10 ms |
| All | $[\text{Ca}^{+}]_{i,\infty}$ | 5e[mol] |
| All | $\underline{g}^{ka}_{A-Type\ K}$ | 0.00015 S/cm$^2$ |
| All | $\underline{g}^{nca}_{Ca-N}$ | 0.0008 S/cm$^2$ |
| All | $\underline{g}^{lca}_{Ca-L}$ | 0.005 S/v |
| All | $\underline{g}^{sk}_{Ca-K}$ | 2e S/cm$^2$ |
| All | $\underline{g}^{BK}_{(Ca,V)-K}$ | 0.0002 S/cm$^2$ |
| All | $E^{ichan2}_{L}$ | -60.06 |
| All | $\underline{g}^{ichan2}_{L}$ | 0.00018 S/cm$^2$ |
| Soma | $\underline{g}^{ichan2}_{Na}$ | 0.12 S/cm$^2$ |
| Soma | $\underline{g}^{ichan2}_{K,fast}$ | 0.013 S/cm$^2$ |
| Soma | $vshift_{ma}$ | 53.75 mV† |
| Soma | $vshift_{mb}$ | 15 mV |
| Soma | $vshift_{ha}$ | 73.75 mV |

| | | |
|---|---|---|
| Soma | $vshift_{hb}$ | 12.5 mV† |
| Soma | $vshift_{nfa}$ | 18 mV |
| Soma | $vshift_{nfb}$ | 43 mV |
| Soma | $vshift_{nsa}$ | 30 mV |
| Soma | $vshift_{nsb}$ | 55 mV |
| Dendrites | $\underline{g}_{Na}^{ichan2}$ | 0.12 S/cm$^2$ |
| Dendrites | $\underline{g}_{Na}^{ichan2}$ | 0 S/cm$^2$ |
| Dendrites | $\underline{g}_{K,fast}^{ichan2}$ | 0.013 S/cm$^2$ |
| Dendrites | $vshift_{ma}$ | 43 mV |
| Dendrites | $vshift_{mb}$ | 15 mV |
| Dendrites | $vshift_{ha}$ | 65 mV |
| Dendrites | $vshift_{hb}$ | 12.5 mV |
| Dendrites | $vshift_{nfa}$ | 18 mV |
| Dendrites | $vshift_{nfb}$ | 43 mV |
| Dendrites | $vshift_{nsa}$ | 30 mV |
| Dendrites | $vshift_{nsb}$ | 55 mV |

*Table 5: Mossy cell biophysics. ** indicates parameters fitted to Howard et al., 2007, † indicates manual adjustment for excitability, to ensure compatibility with iPSC fitted models.*

| Segment(s) | Param | Value |
|---|---|---|
| All | $E_L^{ichan2}$ | -59 |
| All | $\tau_{Ca}^{ccanl}$ | 10 ms |
| All | $[Ca^+]_{i,\infty}$ | $5e^{-6}$ [mol] |
| Dendrites | $\underline{g}_{A-Type\ K}^{ka}$ | 1.650234883413953e-5** S/cm$^2$ |
| Dendrites | $\underline{g}_{Ca-N}^{nca}$ | 3.024649031467018e-5** S/cm$^2$ |
| Dendrites | $\underline{g}_{Ca-L}^{lca}$ | 0.00017469257304634** S/cm$^2$ |
| Dendrites | $\underline{g}_{Ca-K}^{sk}$ | 0.022809187461813662** S/cm$^2$ |
| Dendrites | $\underline{g}_{(Ca,V)-K}^{BK}$ | 0.03039610718265275** S/cm$^2$ |
| Dendrites | $\underline{g}_{hyf}^{hyf}$ | 9.297236447233174e-7** S/cm$^2$ |
| Dendrites | $\underline{g}_{hys}^{hys}$ | 5.951727349497086e-6** S/cm$^2$ |
| Dendrites | $vshift_{ma}$ | 43 mV |
| Dendrites | $vshift_{mb}$ | 15 mV |
| Dendrites | $vshift_{ha}$ | 65 mV |
| Dendrites | $vshift_{hb}$ | 12.5 mV |

| | | |
|---|---|---|
| Dendrites | $vshift_{nfa}$ | 18 mV |
| Dendrites | $vshift_{nfb}$ | 43 mV |
| Dendrites | $vshift_{nsa}$ | 30 mV |
| Dendrites | $vshift_{nsb}$ | 55 mV |
| Soma | $vshift_{ma}$ | 58.5 mV ** † |
| Soma | $vshift_{mb}$ | 27.0368598176269 mV** |
| Soma | $vshift_{ha}$ | 124.5 mV ** † |
| Soma | $vshift_{hb}$ | 18.61654707432046 mV** |
| Soma | $vshift_{nfa}$ | 30.701653087965823 mV** |
| Soma | $vshift_{nfb}$ | 24.156010322453657 mV** |
| Soma | $vshift_{nsa}$ | 44.694885585078325 mV** |
| Soma | $vshift_{nsb}$ | 17.74053364029013 mV** |
| Soma | $\underline{g}_{Na}^{ichan2}$ | 0.055866501682604486 S/cm$^2$ ** |
| Soma | $\underline{g}_{K,fast}^{ichan2}$ | 0.0286296712742244 S/cm$^2$ ** |
| Soma | $\underline{g}_{K,slow}^{ichan2}$ | 0.014713327547369214 S/cm$^2$ ** |
| Soma | $\underline{g}_{L}^{ichan2}$ | 8.0500055186908e-06 S/cm$^2$ ** |
| Soma | $\underline{g}_{A-Type\ K}^{ka}$ | 0.000015360279879 S/cm$^2$** |

| | | |
|---|---|---|
| Soma | $\underline{g}_{Ca-N}^{nca}$ | 0.000046076963887 S/cm$^2$** |
| Soma | $\underline{g}_{Ca-L}^{lca}$ | 0.00076494065621 S/cm$^2$** |
| Soma | $\underline{g}_{Ca-K}^{sk}$ | 0.021562445425285 S/cm$^2$** |
| Soma | $\underline{g}_{(Ca,V)-K}^{BK}$ | 0.030179002060815 S/cm$^2$** |
| Soma | $\underline{g}_{hyf}^{hyf}$ | 0.000009836462022 S/cm$^2$** |
| Soma | $\underline{g}_{hys}^{hys}$ | 0.000008197063742 S/cm$^2$** |
| Dendrites | $\underline{g}_{L}^{ichan2}$ | 4.4e$^{-5}$ S/cm$^2$ |
| Proximal Dendrites | $\underline{g}_{Na}^{ichan2}$ | 0.12 S/cm$^2$ |
| Distal Dendrites | $\underline{g}_{Na}^{ichan2}$ | 0.0 S/cm$^2$ |
| Proximal Dendrites | $\underline{g}_{K,fast}^{ichan2}$ | 0.0005 S/cm$^2$ |
| Distal Dendrites | $\underline{g}_{K,fast}^{ichan2}$ | 0.0 S/cm$^2$ |

*Table 6: HIPP cell biophysics. † indicates manual adjustment for excitability*

| **Segment(s)** | **Param** | **Value** |
|---|---|---|
| Dendrites | $\underline{g}_{L}^{ichan2}$ | 3.6e$^{-5}$ S/cm$^2$ |
| All | $E_{L}^{ichan2}$ | -70.45 |
| All | $\tau_{Ca}^{ccanl}$ | 10 ms |
| All | $[Ca^{+}]_{i,\infty}$ | 5e$^{-6}$ [mol] |

| | | |
|---|---|---|
| Dendrites | $\underline{g}^{ka}_{A-Type\ K}$ | 0.0008 S/cm$^2$ |
| Dendrites | $\underline{g}^{nca}_{Ca-N}$ | 0.0 S/cm$^2$ |
| Dendrites | $\underline{g}^{lca}_{Ca-L}$ | 0.0015 S/cm$^2$ |
| Dendrites | $\underline{g}^{sk}_{Ca-K}$ | 0.003 S/cm$^2$ |
| Dendrites | $\underline{g}^{BK}_{(Ca,V)-K}$ | 0.003 S/cm$^2$ |
| Dendrites | $\underline{g}^{hyf}_{hyf}$ | 1.5e$^{-5}$ S/cm$^2$ |
| Dendrites | $\underline{g}^{hys}_{hys}$ | 1.5e$^{-5}$ S/cm$^2$ |
| Soma | $vshift_{ma}$ | 58 mV † |
| Soma | $vshift_{mb}$ | 15 mV |
| Soma | $vshift_{ha}$ | 82 mV † |
| Soma | $vshift_{hb}$ | 12.5 mV |
| Soma | $vshift_{nfa}$ | 18 mV |
| Soma | $vshift_{nfb}$ | 43 mV |
| Soma | $vshift_{nsa}$ | 30 mV |
| Soma | $vshift_{nsb}$ | 55 mV |
| Soma | $\underline{g}^{ichan2}_{L}$ | 3.6e$^{-5}$ S/cm$^2$ |
| Soma | $\underline{g}^{ichan2}_{Na}$ | 0.2 S/cm$^2$ |

| | | |
|---|---|---|
| Soma | $\underline{g}^{ichan2}_{K,fast}$ | 0.006 S/cm$^2$ |
| Soma | $\underline{g}^{ichan2}_{K,slow}$ | 0.001 S/cm$^2$ |
| Soma | $\underline{g}^{ka}_{A-Type\ K}$ | 0.0008 S/cm$^2$ |
| Soma | $\underline{g}^{nca}_{Ca-N}$ | 0.0008 S/cm$^2$ |
| Soma | $\underline{g}^{lca}_{Ca-L}$ | 0.0015 S/cm$^2$ |
| Soma | $\underline{g}^{hyf}_{hyf}$ | 0.000015 S/cm$^2$ |
| Soma | $\underline{g}^{hys}_{hys}$ | 0.000015 S/cm$^2$ |
| Soma | $\underline{g}^{sk}_{Ca-K}$ | 0.003 S/cm$^2$ |
| Dendrite | $\underline{g}^{ichan2}_{Na}$ | 0.2 S/cm$^2$ |
| Dendrite | $\underline{g}^{ichan2}_{K,fast}$ | 0.006 S/cm$^2$ |
| Dendrite | $\underline{g}^{ichan2}_{Na}$ | 0 S/cm$^2$ |
| Dendrite | $\underline{g}^{ichan2}_{K,fast}$ | 0 S/cm$^2$ |

*3.3. Network connectivity and other biophysical parameters*

*Table 7: Connectivity parameters for biophysical model*

| **From** | **To** | **Sections** | **Connectivity** | **Kind** | **Mechanism** | **Delay** | **Weight** |
|---|---|---|---|---|---|---|---|
| PP | GC | Dendrite | 0.2 | C | gcdendampa | 3 | 0.002 |
| PP | BC | Apical Dendrite | 0.2 | C | bcddendaampa | 3 | 0.001 |

| | | | | | | | |
|---|---|---|---|---|---|---|---|
| GC | BC | adend | 0.167 | D | bcpdendampa | 0.8 | 0.0141 |
| GC | MC | adend | 0.067 | D | mcpdendampa | 1.5 | 0.0002 |
| GC | HIPP | adend | 0.5 | D | hipppdendampa | 1.5 | 0.005 |
| BC | GC | soma | 0.2 | D | gcsomagaba | 0.85 | 0.0048 |
| BC | BC | Apical Dendrite | 0.333 | D | bcbcmdendgaba | 0.8 | 0.0076 |
| BC | MC | soma | 0.2 | D | mcsomagaba | 1.5 | 0.0015 |
| MC | GC | pdend | 0.4 | D | gcdendampa | 3 | 0.0003 |
| MC | BC | Apical Dendrite | 0.167 | D | bcmdendaampa | 3 | 0.0003 |
| MC | MC | adend | 0.2 | D | mcmcpdendampa | 2 | 0.0005 |
| MC | HIPP | bdend | 0.333 | D | hippmdendampa | 3 | 0.0002 |
| HIPP | GC | Dendrite | 0.32 | D | gcdendgaba | 1.6 | 0.0005 |
| HIPP | BC | Dendrite | 0.667 | D | hcbcmdendgaba | 1.6 | 0.0005 |
| HIPP | MC | cdend | 0.267 | D | mcpdendgaba | 1 | 0.0015 |

*Table 8: Synapse parameters for biophysical model*

| **Type** | **Name** | **Mechanism** | $\tau_1$ | $\tau_2$ | **e** |
|---|---|---|---|---|---|

| | | | | | |
|---|---|---|---|---|---|
| GABA | gcsomagaba | Exp2Syn | 0.26 | 5.5 | -70 |
| GABA | gcdendgaba | Exp2Syn | 0.5 | 6 | -70 |
| GABA | bcbcmdendgaba | Exp2Syn | 0.16 | 1.8 | -70 |
| GABA | hcbcmdendgaba | Exp2Syn | 0.4 | 5.8 | -70 |
| GABA | mcsomagaba | Exp2Syn | 0.3 | 3.3 | -70 |
| GABA | mcpdendgaba | Exp2Syn | 0.5 | 6 | -70 |
| AMPA | gcdendampa | Exp2Syn | 1.5 | 5.5 | 0 |
| AMPA | bcddendaampa | Exp2Syn | 2 | 6.3 | 0 |
| AMPA | bcpdendampa | Exp2Syn | 0.3 | 0.6 | 0 |
| AMPA | bcmdendaampa | Exp2Syn | 0.9 | 3.6 | 0 |
| AMPA | mcddendampa | Exp2Syn | 1.5 | 5.5 | 0 |
| AMPA | mcpdendampa | Exp2Syn | 0.5 | 6.2 | 0 |
| AMPA | mcmcpdendampa | Exp2Syn | 0.45 | 2.2 | 0 |
| AMPA | hipppdendampa | Exp2Syn | 0.3 | 0.6 | 0 |
| AMPA | hippmdendampa | Exp2Syn | 0.9 | 3.6 | 0 |

*Table 9: Geometry of cells within biophysical model*

| **Cell** | **Section** | **Property** | **Value** |
|---|---|---|---|
| GC | All | $R_a$ | 184 Ω cm |
| GC | Soma | Diameter | 16.8 μm |
| GC | Soma | *L* | 16.8 μm |
| GC | Soma | $C_m$ | 1 μF/cm$^2$ |
| GC | Dendrites$_{GCL,P}$ | $C_m$ | 1 μF/cm$^2$ |
| GC | Dendrites$_{M,D}$ | $C_m$ | 1.6 μF/cm$^2$ |
| GC | Dendrites$_{GCL}$ | *L* | 50 μm |
| GC | Dendrites $_{P,M,D}$ | *L* | 150 μm |
| GC | Dendrites | Diameter | 3 μm |
| BC | All | $C_m$ | 1.4 μF/cm$^2$ |
| BC | All | $R_a$ | 100 Ω cm |
| BC | Soma | *L* | 20 μm |
| BC | Soma | Diameter | 15 μm |
| BC | Apical Dendrites | *L* | 75 μm |
| BC | Basal Dendrites | *L* | 50 μm |
| BC | Dendrite$_A$ | Diameter | 4 μm |
| BC | Dendrite$_B$ | Diameter | 3 μm |
| BC | Dendrite$_C$ | Diameter | 2 μm |
| BC | Dendrite$_D$ | Diameter | 1 μm |
| MC | All | $R_a$ | 100 Ω cm |
| MC | Soma | *L* | 20 μm |
| MC | Soma | Diameter | 20 μm |
| MC | Soma | $C_m$ | 0.6 μF/cm$^2$ |
| MC | Dendrites | $C_m$ | 2.4 μF/cm$^2$ |

| | | | |
|---|---|---|---|
| MC | Dendrites | *L* | 50 µm |
| MC | $Dendrite_A$ | Diameter | 5.78 µm |
| MC | $Dendrite_B$ | Diameter | 4 µm |
| MC | $Dendrite_C$ | Diameter | 2.5 µm |
| MC | $Dendrite_D$ | Diameter | 1 µm |
| HIPP | All | $R_a$ | 100 Ω cm |
| HIPP | Soma | *L* | 20 µm |
| HIPP | Soma | Diameter | 10 µm |
| HIPP | All | $C_m$ | 1.1 µF/cm$^2$ |
| HIPP | Apical Dendrites | *L* | 75 µm |
| HIPP | Basal Dendrites | *L* | 50 µm |
| HIPP | $Dendrite_A$ | Diameter | 3 µm |
| HIPP | $Dendrite_B$ | Diameter | 2 µm |
| HIPP | $Dendrite_C$ | Diameter | 1 µm |

### 4. Additional Discussion

#### *4.1. Effects of lithium on healthy control WTA dynamics: schematic of theory*

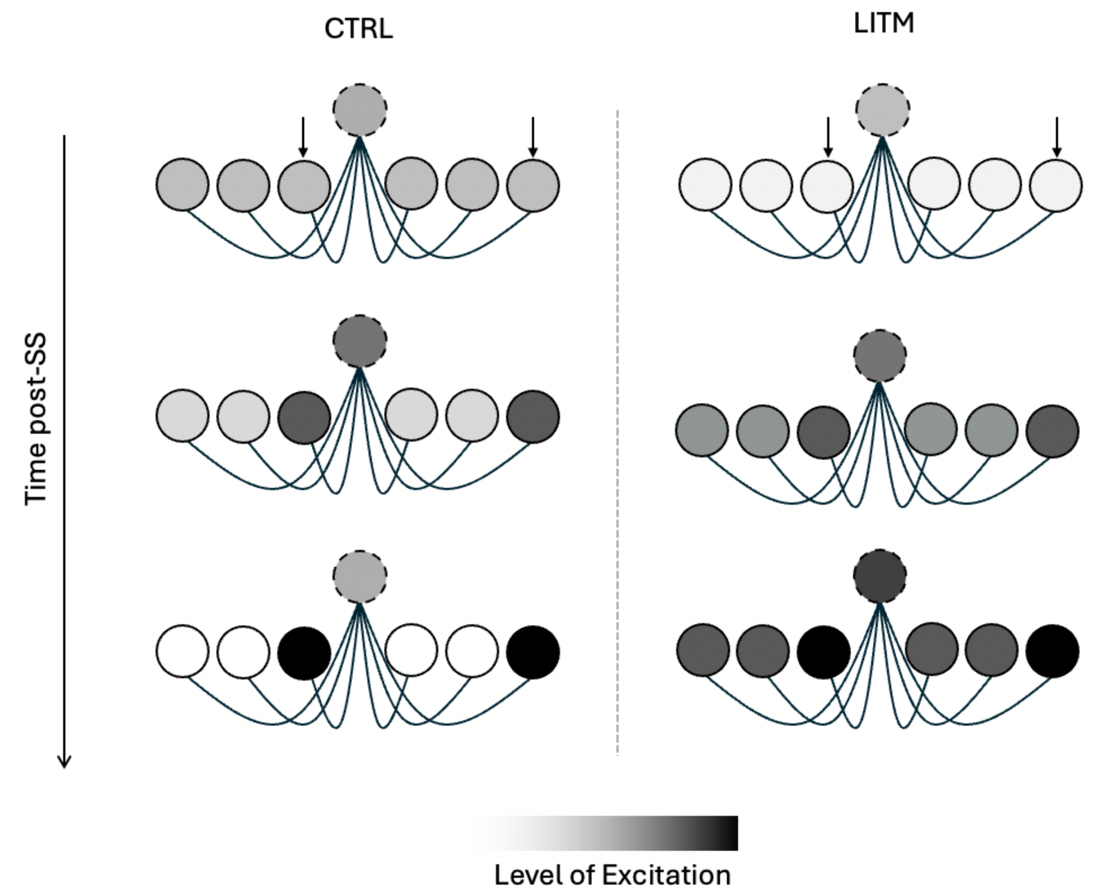

*Figure 5: Schematic representation of lithium's impacts on WTA dynamics in HCs. Circles with dashed outlines are BCs, while circles with solid lines are GCs. Short arrows pointing towards select GCs indicate those that are spontaneously active. In control conditions, spontaneous activity will lead to the depolarization of those select GCs, depolarizing the one BC. That BC in turn will inhibit all neurons. The neurons with the highest activity levels will "escape" from this inhibition, while the activity in the remaining GCs will reduce. Lithium exposure reduces GC sensitivity to both excitatory and inhibitory currents. Therefore the widespread BC inhibition caused by initial GC depolarisation is not as effective at inhibiting the remaining GCs, leading to a more "relaxed" GC competition.*

*4.2. Degeneracy*

Model fitting procedures often resulted in multiple families of parameters that produce the same behaviour: a phenomenon known as parameter degeneracy[13]. It is therefore likely that starting the optimisation procedure with different initial conditions may produce different parameter sets that, once implemented in a cellular model, will generate comparable behaviour to the models used in the present study. As a sanity check, we ran our parameter optimisation procedure initialised with different random seeds, and found that the parameter sets were relatively similar to one another, minimising concerns about degeneracy. This may not be the case if a different optimisation algorithm was used, however.

Degeneracy is not necessarily a complete limitation. There is evidence in biological networks that channel properties and cell morphology are highly heterogeneous, but may produce the same measured electrophysiological behaviour[14–16]. Degeneracy, or redundancy, may confer an advantage over having a homogeneous population, as it may mitigate the effects of a potential failure in one mechanistic system, as there are other systems to rely on. To assess degrees of heterogeneity in cultured iPSC neurons derived from individuals with BD, an unbiased in vitro classification using electrophysiological and morphological cell properties may be beneficial, following similar classification work done on cultured DG interneurons[17]. This type of classification work would help modellers decide whether cellular heterogeneity should be incorporated into their network models, which would have further implications for neural computation[18].
